# Comparative performance of reference-based metagenomic tools to identify species-level taxa among families of bacteria: benchmarking *Mycobacteriaceae* and *Neisseriaceae*

**DOI:** 10.1101/2025.06.23.661092

**Authors:** Luke B. Harrison, Juba O. Ahmed, Gnéré Coulibaly, Frédéric J. Veyrier

## Abstract

Hypotheses concerning the ecology and evolution of bacteria commonly relate to the presence and abundance of species in various settings and conditions. Shotgun metagenomics may address these hypotheses, which previously relied on PCR or culture. However, the problem of determining the presence/absence of a given species of interest is not trivial, particularly when closely related species are present in the reference database or metagenomic sample. Reference-based methods to detect species-level taxa mostly rely on thresholding of aligned reads or mapped k-mers or derivative metrics like genomic coverage, and create a trade-off between recall/completeness and precision/purity. New methods for species-level profiling (YACHT, metapresence and sylph) have recently been published. Here we test the performance of these methods, along with Kraken2/bracken and MetaPhlAn4, to detect related species of interest using simulated metagenomic samples from genomes in the families *Mycobacteriaceae* and *Neisseriaceae*, which contain closely related genomes. Among methods tested, metapresence, when used with an alignment quality filter, and sylph offer the best overall performance. Sylph maintains high precision but requires a depth of coverage greater than approximately 0.1x to reliably detect a genome’s presence. Metapresence has a lower limit of detection of hundreds of reads but this is balanced against relatively lower precision. Both methods are relatively robust to the presence of reads from genomes outside the groups of interest. We demonstrate the application of these methods in two real-world datasets: a mycobacterial community in a drinking water system and the community of *Neisseriaceae* present in the human oral cavity.

**Importance:** Detecting which bacterial species of interest are present in a given sample is fundamental to studies of microbial ecology and evolution, and to applied microbiology (e.g. clinical diagnostics). Culture-dependent and independent (e.g. PCR) approaches are increasingly complemented by metagenomic approaches, but methods to accurately identify specific low-abundance species-level genomes in a shotgun metagenomic sample are still being refined. Here we comprehensively test YACHT, Kraken2/bracken, metapresence, MetaPhlAn4, and sylph using two simulated datasets of bacterial families, *Mycobacteriaceae* and *Neisseriaceae* that contain closely related species. Our simulations exploit natural genomic diversity to create a challenging benchmark. We demonstrate that metapresence and sylph perform best, with the former being well-suited to low-biomass host-associated datasets, and the latter with environmental metagenomic samples. This study is the first extensive benchmark of these methods for this use case, and demonstrates these methods can accurately identify closely related species of interest.

## Introduction

Determining the presence or absence of a given species in an ecological niche can be accomplished through culture-dependent and culture-independent methods, frequently based on PCR amplification of marker genes with amplicon sequencing, for example of 16S rRNA genes, or *hsp65* mycobacteria-specific PCR^1^. Shotgun metagenomic sequencing has the potential to characterize the composition of microbial communities and identify the presence of bacterial species, irrespective of cultivability, and to concomitantly offer insight into the aggregate functional content of the microbial genomes^2,3^. Conceptually, these approaches can broadly be divided into reference-free methods that seek to produce metagenome-assembled genomes (MAGs) from de novo assembly of sequencing reads^3,4^, and reference-based methods that rely on a database of genomes (which may also be MAGs) against which sequencing reads are mapped^2^. Most recently, with new reference-based profiling algorithms, shotgun metagenomics offers the promise of accurate species-level identification in complex metagenomic samples, at low abundance^5–8^. This opens the door to the broad application of metagenomic methods as tools for the targeted identification of specific species-level taxa, among an arbitrary taxonomic group of interest, in both existing and prospectively collected samples without employing resource-intensive upstream wet-lab workflows (e.g. specific PCR with amplicon sequencing or bait-capture).

Identifying specific species present is relevant to test hypotheses concerning the ecology and evolution of microorganisms, and to interrogate the etiological agents of infectious diseases. The family *Mycobacteriaceae* is a clinically relevant group of mostly environmental bacteria, whose precise niches are informed by both culture-dependent and culture-independent^9–11^ methods, for which many questions remain^12^. More precise delineation of the habitats of mycobacteria may inform targeted interventions to reduce exposures and disease among vulnerable populations^12,13^. A second example is the tropism of species of the family *Neisseriaceae*, whose closely related members inhabit narrow symbiotic niches amongst their vertebrate hosts^14^, and whose evolutionary history is driven in part by niche-specific host interactions^15^. Previously, to address these hypotheses, researchers investigating these groups have employed techniques including cultures using enriched, selective and/or differential media, targeted PCRs and amplicon sequencing^16^, mass spectrometry^17^, and some metagenomics approaches^10,14^.

Interrogating these questions using metagenomic sequencing, where testing hypotheses requires the discrimination of the presence and abundance of defined organisms at the species level, at a potentially very low abundance, is best approached using a reference-based method^6^. This metagenomic problem differs from the more general problem of taxonomic classification of sequencing reads or community profiling (e.g. microbiome composition), and we focus on the performance of these methods to determine the presence or absence of a set of (mostly) species-level genomes in a given sample, irrespective of overall community composition.

However, this is not trivial^6^ because bacteria share significant genomic sequence similarity through vertical evolution (e.g. phylogenetic relatedness) or horizontal evolution (e.g. gene transfer)^5,6,18^. For these reasons, and because bacterial species may be named and defined for historical or phenotypic reasons, despite having high average nucleotide identity (ANI), individual sequencing reads may map well to the genomes of multiple species^5,6,19^. Furthermore, within a species, those organisms with open pangenomes may harbor significant variation in gene content relative to a given reference genome^19^. This complicates both the assignment of reads to a given taxon and the determination of which species are truly present/absent in a given metagenome^19^. Methods based on k-mer matching (e.g. Kraken2^21^ and related algorithms) are known to yield false positive read classifications at the species level, necessitating strict thresholding, and whose precision at the species level has been reported as low^5^. These problems may be further compounded in host-associated metagenomic studies due to low biomass, and/or incomplete host sequence removal^22^.

These problems have been recognized, and several algorithms to improve the recall (ability to detect a genome that is truly present, also called sensitivity or completeness) and precision (the fraction of genomes identified by the method as present that are truly present and not false positives, also called purity) of metagenomic tools have been published. These incorporate information such as spatial mapped read distributions, and read sequence divergence (e.g. KMCP^7^) or employ marker gene approaches (e.g. MetaPhlAn4^23^). More recently, several new algorithms for arbitrary database reference-based, species-level metagenomic profiling have been published. Among them, two are k-mer-based tools that use containment ANI (YACHT^8^ and sylph^5^) and one is read alignment distribution-based (metapresence^6^). Here, we test the performance of these tools for the specific problem of determining which related species within a given taxonomic group are present within a metagenomic sample, regardless of the overall community composition. We explored performance of these methods on simulated microbial communities of closely related species in a single family, choosing taxa pragmatically (here *Mycobacteriaceae* and *Neisseriaceae*), that may contain species and species complexes of variable genetic interrelatedness. In an attempt to create a realistic, but challenging real-world benchmark, these datasets are expected to contain species with open pangenomes and significant HGT (in the case of naturally competent *Neisseriaceae*, HGT is particularly widespread^24^) where pairwise ANI is variable and may exceed 95%. We further enriched our simulation with natural genome evolutionary dynamics by exploiting the diversity of publicly available genome sequences to create our simulated datasets rather than relying only on reference genome sequences themselves. We then compared these methods against Kraken2/bracken, a widely used k-mer-based method and MetaPhlAn4^23^, a marker-gene-based approach.

These methods implicitly or explicitly use thresholds to define a given genome’s presence: this creates a trade-off between the ability to detect a genome that is truly present (measured using recall) and the incorrect detection of genomes not truly present (measured using precision). We therefore compare the performance of these methods, as well as simpler methods based on aligned read count and breadth of coverage-based thresholds, using precision-recall curves (PRC). We further tested the effect of potentially confounding reads from incomplete host genome removal, and in the presence of reads derived from mock communities of microorganisms using both a GTDB-based simulation and the CAMI II challenge datasets^24^. We also investigated their sensitivity to incomplete reference databases, and their recall performance at low abundance. Finally, we present case-studies of the application of these methods to the profiling of mycobacterial communities present in a water distribution system^10^ and *Neisseriaceae* present in the human oral cavity^14,25,26^.

## Materials and Methods

### Reference genome databases and simulated datasets

We assembled a reference genome database for the family *Mycobacteriaceae* (*n=*252) and used that of Chenal et al.^27^ for *Neisseriaceae* (*n*=70), both using existing sequence data (NCBI genome accessions in Tables S1 and S2). These databases were curated to select the highest quality genome for each species, preserving named species even if they shared significant genomic similarity but have important phenotypic distinctions (e.g. *Neisseria meningitidis* and *N. gonorrhoeae*, ANI = 95%; *Mycobacterium tuberculosis* and *M. canettii*, ANI = 99.2%). We also retained named subspecies of significance for *Mycobacteriaceae* (e.g. *M. abscessus* subsp. *bolletii*, *massiliense*, and *abscessus,* ANI ∼97%). For *Mycobacteriaceae*, a core genome phylogeny was estimated by first using roary^28^ to produce a core genome alignment using the NCBI GFF3 gene annotation files (roary parameters-e mafft-i 60-cd 99), followed by phylogenetic reconstruction using iqtree v2.2.2.7^29^. To illustrate genome relatedness, pairwise average nucleotide identity (ANI) was calculated using fastANI^30^. The same procedure was performed for the *Neisseriaceae* dataset using fastANI and the phylogeny estimated by Chenal et al.^27^.

We then simulated 500 replicate shotgun metagenomes for each database (simulation procedure summarized in Fig. S1), consisting of paired-end Illumina sequencing reads simulated from a randomly selected subset of species present in our reference databases (variable *n* species per replicate, drawn from normal distributions with means of 15 and 30, respectively and a standard deviation of 4). To represent the intraspecific genomic variability present in natural species, including variation in gene content due to horizontal gene transfer and other processes: for each named species’ reference genome, we randomly retrieved a corresponding genome sequence from the same named species from NCBI (RefSeq, all genomes assigned to *Mycobacteriaceae or Neisseriaceae*, retrieved using NCBI datasets, 2025/02/11, Supplementary Tables S3, and S4). Using a custom script (Supplementary materials, Zenodo archive, using the software packages fastANI 1.32^30^, seqkit v.2.3.1^31^, and samtools v.1.17^32^, ChatGPT o4-mini-high was used as a coding aid to refine and troubleshoot scripts used in this manuscript, but not to conceive or create them), we verified the taxonomic identity of the chosen genome by comparing its ANI to our reference genome database and ensuring the reference genome for the (sub)species being simulated was the top ANI hit to the randomly selected genome, and had ANI > 95%. In the case where no suitable alternative genome was available for a given taxon or it was an unnamed species, the reference genome was used for simulations. We then simulated paired-end reads using Mason^33^ and first used a read length of 100bp, as this should present the greatest challenge, and then repeated selected analyses using a read length of 150bp, to achieve a mean depth drawn from an exponential distribution with a mean randomly selected from values between 0.01 and 10x (Fig. S1). We then injected both simulated sequencing error and single nucleotide polymorphism (SNPs, at a rate randomly selected from 0.02, 0.01, 0.001, or 0.0001 substitutions per site), all while assuring an ANI of no less than 95% from the reference genome for the selected species was maintained in the final mutated genome used to simulate sequencing reads. Finally, simulated reads from each selected species were combined into a synthetic metagenome.

### Execution of metagenomic tools and alignment-based methods

To add context to the results, we also defined simple threshold-based species-level genome identification tools using read alignments. Reads were first competitively aligned to the respective concatenated reference database using bowtie2 (options –very-sensitive --end-to-end)^34^, and then filtered for alignment mapping quality (none, MAPQ2, 3, 5, 10, 20 and 30) using samtools. Read count and breadth of coverage-based methods were implemented using scripts to process the results of samtools’ coverage tool, and report a species present if the aligned read count or breadth of genomic coverage (defined as fraction of target bases covered by at least one aligned read^6,35^) exceeded a predefined threshold. Metapresence^6^ is an alignment-based method, so aligned reads contained in a BAM file, at varying mapping quality filter levels, were used to execute the metapresence script. YACHT^8^ and sylph^5^ were executed after building the respective reference k-mer sketches using default parameters. For each method, thresholds were varied among the relevant ranges. For metapresence, the thresholding parameters of the Fraction of Unexpected Gaps (FUG) and Breadth-Expected breadth Ratio (BER)^6^ were varied along their ranges of 0-1. For YACHT, there are two main thresholding parameters: the ANI cutoff at which genomes are considered the same species, and the minimum exclusive breadth of coverage. Executions were performed at varying thresholds for each. Sylph can also be tuned for ANI threshold, which was allowed to vary between 0.95 and 0.99. Sylph has three other tuning parameters:-c, for the fraction of k-mers sampled for sketching of both genomes and reads, --min-count-correct, the k-mer multiplicity required for coverage correction, and --min-number-kmers which serve as a minimum threshold to output results. Our tests demonstrated essentially no meaningful impact of these parameters in this analysis, yielding the same mean precision and recall at ANI = 0.95 across replicates, so these were kept at the defaults. Kraken2/bracken^21^ was executed using default parameters, first with using a custom index consisting only of our reference genome databases and then with the “Standard” precompiled index (https://benlangmead.github.io/aws-indexes/k2, Oct 15th, 2025) combined with our reference databases. MetaPhlAn4^23^ was executed using default parameters. In both cases, these algorithms consider only species-level taxa and use the NCBI taxonomy database. To evaluate their performance against the above methods, we translated our taxonomy to NCBI’s and collapsed the analysis to species-level for these methods only (correspondence in Tables S1 and S2).

### Precision-recall curves

The relative performance of these methods to recover the genomes that were used to generate the simulated datasets was compared using precision-recall curves (PRC) after calculating the relevant statistics (true positives, false positives, false negatives, and true negatives) across the relevant parameter/threshold spaces for each method in each replicate metagenome. Results are plotted for each threshold in precision-recall space at the mean of each metric across replicates (± standard deviation) using a python script (Supplementary materials, Zenodo archive) and the following packages: numpy^36^, pandas^37^, Scikit-learn^38^ and matplotlib^39^. PR curves were compared using area under the curve measurements, and optimal parameters were determined by selecting the set of parameters that maximized the F_1_-score.

### Recall in relation to simulated genome depth and abundance estimation

A further analysis was conducted to measure recall as a function of the simulated genome depth, binning genomes by depth across all simulated metagenomes for each reference dataset using an R script (Supplementary materials, Zenodo archive). The read count-based threshold method was not tested as it was similar to the breadth-based method but performed less well (see results). For all methods tested, the best performing threshold parameters from the PRC analysis were used for each dataset and alignment filtering stringency (Table 1). This analysis was also repeated binning genomes across the actual simulated read counts. Recall was summarized using the mean recall in a given bin, using a bootstrap resampling procedure with 1000 replicates to provide a measure of significance (95% bootstrapped range of means). Accuracy of abundance estimation was compared between methods by comparing the true simulated abundance to the predicted abundance by computing the normalized L_1_ (Manhattan distance) metric.

**Table 1.**
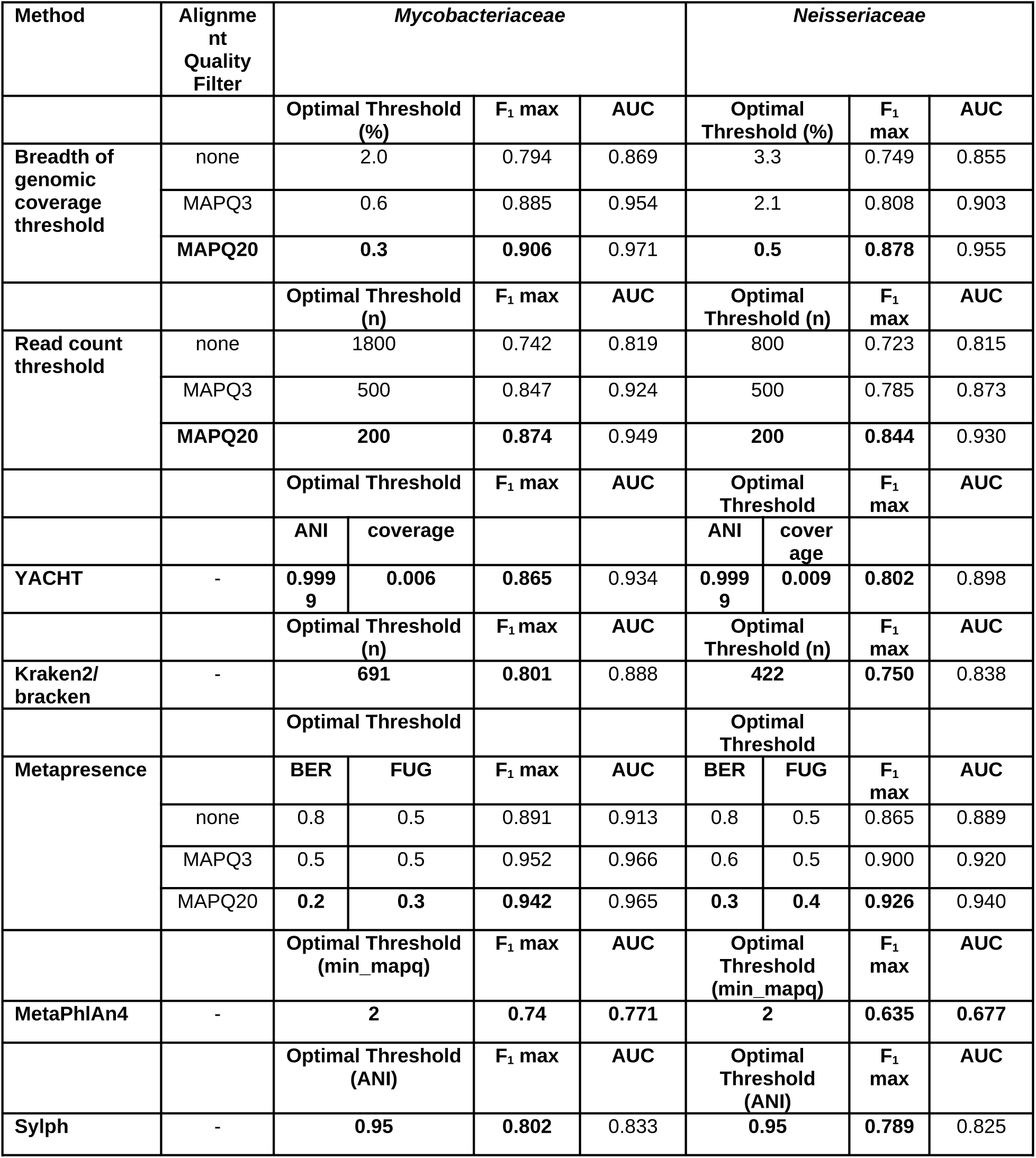
Performance of methods to identify genomes present in a metagenomic sample based on 500 simulated metagenomic datasets (100bp read length) of species of *Mycobacteriaceae* and *Neisseriaceae*. Optimized threshold parameters are determined by maximizing the F_1_-score. For alignment-based methods, results for mapping quality filtering at MAPQ3 and MAPQ20 are shown.

### Precision in relation to incomplete host removal and the presence of other microbial reads

To assess the robustness of these methods to incomplete removal of host reads, we first performed an experiment using reads simulated from the human genome (NCBI Genome accession GCF_000001405.40). Using mason^33^, we simulated Illumina reads from the human genome to an average depth of 1.0x. We repeated this 10 times then combined these reads with a randomly selected sample of 10 of our simulated metagenome replicates from the *Mycobacteriaceae* and *Neisseriaceae* databases and repeated the execution of the methods, comparing precision between the 10 simulated metagenomes, with and without additional human host reads.

To determine whether the inclusion of other microbial sequences not reflected in the reference genome impacts precision, we first simulated reads from microbial genomes drawn from release 220 (r220) of the Genome Taxonomy Database (GTDB)^40^. We removed genomes attributed to our families of interest in the GTDB taxonomy, and then randomly selected 100 genomes from the GTDB r220, simulated reads using mason, injecting both read and SNP errors, at a random depth, as above, but without resampling genomes from NCBI. We repeated this 100 times, and concatenated these synthetic GTDB metagenomes with random samples of 100 each of the *Mycobacteriaceae* and *Neisseriaceae* simulated metagenomes (simulated at read length 100). We then compared precision of the methods with and without the additional microbial reads. We then repeated a similar experiment using the mock communities in the CAMI II challenge datasets^24^. We randomly sampled mock metagenomes from the CAMI II Marine and Plant Rhizosphere datasets (with replacement) and fused these with our *Mycobacteriaceae* metagenomes. For *Neisseriaceae*, we combined our simulated reads with randomly sampled metagenomes from the CAMI II Toy Human Microbiome Project dataset^24^. Because the CAMI II datasets were generated with 150bp read lengths, we fused CAMI II reads with our *Mycobacteriaceae* and *Neisseriaceae* datasets simulated at that read length. In the case where these read sets already contained reads generated from species-level taxa present in our reference, we merged the relevant species into our truthsets for each given replicate.

The degree to which the precision of selected methods is affected by an incomplete reference database was assessed using a database subsampling experiment. For each reference database, a set of subsampled reference databases was created by randomly removing an increasing fraction of reference genomes, in steps of 10% of the total reference database size, generating 20 replicates of the subsampled reference genome database for each level of genome removal using a python script (Supplementary materials). Then, these sets of subsampled databases were used with the various methods to estimate precision at each level of genome removal using a randomly selected sample of 20 simulated metagenomes per family from the initial 500 metagenomes for each family, such that the samples may contain reads from species no longer present in the subsampled reference database.

### Case studies

To illustrate the best performing methods’ performance on real-world datasets, we retrieved two shotgun metagenomic datasets. For *Mycobacteriaceae*, we downloaded short-read metagenomes (NCBI Bioproject PRJNA1081894) derived from an interrogation of a chloraminated wastewater system dataset^10^, for which both culture and MAG-based approaches were employed in the original article. The MAG approach included read filtering, pooling across samples, co-assembly with metaSPAdes^41^, followed by genome binning using DAS Tools^42^.

MAGs were only retained if they were high-quality with completeness > 50%. Presence of MAGs in each sample were then determined with coverM^10^. Here, adapters were trimmed from the raw reads using fastp v0.240^43^ and aligned against the *Mycobacteriaceae* reference database using bowtie2^33^. Sylph and metapresence were executed as above, using optimized BER/FUG parameters (Table 1) for metapresence. For *Neisseriaceae*, we downloaded oral cavity metagenomes from the dorsum of tongue and supragingival plaque of healthy humans from Phase I of the human microbiome project (HMP, SRA sample accessions in Table S5)^25^, ensuring samples for each site were from different individuals. HMP Phase I reads already have host (human) reads removed, and sequencing adapters trimmed. These reads were aligned to the *Neisseriaceae* database using bowtie2, with sylph and metapresence executed as above.

For both datasets, we extracted the presence/absence and relative overall abundance from the outputs of sylph (using the-u option) and metapresence (obtaining absolute read count from the metapresence output), and scaled the abundances using the corresponding genome length and overall read count to obtain read per kilobase-million (RPKM). For illustrative purposes and comparability, as genome breadth is not directly reported by sylph, we calculated genome breadth for all genomes in the respective reference databases using the unfiltered bowtie2 read alignment and the samtools coverage tool.

## Results

### Datasets

We first assembled reference genome databases for *Mycobacteriaceae* and *Neisseriaceae*. The assembled reference genome databases demonstrated a range of genome relatedness (Fig. S2,S3). The *Mycobacteriaceae* database contained several clades of named species sharing high ANI (e.g. *M. ulcerans*, *M. marinum*, and related species). The core genome phylogeny calculated here (Fig. S2) was similar to recently published analyses^44–46^ and we have opted to use the traditional nomenclature of the family, with a single genus, *Mycobacterium*^47^. The *Neisseriaceae* reference database was notable both for the relatively high ANI within certain clades (e.g. *N. meningitidis*, *N. gonorrhoeae* and other related species of nasopharyngeal *Neisseria*, pairwise ANI ∼94-95%) and for significant horizontal gene transfer leading to unexpectedly high ANI at odds with the core genome phylogeny (Fig. S3). For example, although relatively basal among *Neisseria* in the core genome tree, the human oral commensal *N. elongata* had relatively higher ANI relative to other human oral commensal *Neisseria*, suggesting HGT between these co-occurring species^15^. The overall mean±SD pairwise ANI in these family-level reference genome databases was 0.80±0.02 for *Mycobacteriaceae* and 0.81±0.06 for *Neisseriaceae*. For the 100bp read length simulations, the mean±SD number of genomes per replicate was 29.2±3.7 and 14.8±3.8, respectively, for the *Mycobacteriaceae* and *Neisseriaceae* datasets. Median and interquartile range (IQR) for depth of coverage, simulated read count, and breadth of genomic coverage (fraction of genome covered by at least one read^6,35^) across replicate simulated metagenomes was 0.23x (1.21), 6547 reads (34368), and 20% (68) for *Mycobacteriaceae* and 0.22x (1.17), 2539 reads (13928), and 19% (66) for *Neisseriaceae*. The mean final ANI between the selected, mutated genomes used to simulate reads (Fig. S1) and their corresponding reference genomes was 98.7±1.1 and 98.1±1.5, respectively.

### Precision and recall performance

We next assessed the performance of the methods in order to examine their explicit and implicit trade-offs between recall (the ability of a given method to detect the presence of a given species that is truly present, also called true positive rate, sensitivity or completeness) and precision (the fraction of species identified as present by the method that are truly present in the metagenomic sample, also called purity). These trade-offs were examined visually using precision-recall curves (PRCs, Fig. 1,2, Fig. S4-S7), and by comparing area under the curve (AUC) values for each method (100bp read length: Table 1, 150bp read length: Table S6). The simple breadth-based threshold with alignment filtering was the most optimal classifier tested by AUC criteria (Table 1, S6), and performed better than read-count based thresholds (Fig. 1a,b, 2a,b, S4a,b, S5a,b). PRCs also demonstrated the importance of filtering aligned reads by mapping quality, which yielded better overall classifier performance for the aligned read and breadth threshold-based metrics (Fig. 1a,b, 2a,b, S4a,b, S5a,b, Table 1, S6). However, the alignment threshold-based approaches (breadth, read count) began to sacrifice precision if the threshold was set to achieve recall greater than 0.6-0.7 in these datasets. YACHT performed less well at lower ANI thresholds (Fig. 1c,2c, S4c, S5c, Table 1, S6). Kraken2/bracken performed similarly to unfiltered alignment-based approaches (Fig. 1d, 2d, S4d, S4d, Table 1, S6). Metapresence performed well across both datasets (Fig. 1e,f, 2e,f, S4e,f, S5e,f), and provided very good recall before sacrificing precision, particularly when the input alignment is filtered for mapping quality (Fig. 1f, 2f, Fig. S6, S7). MetaPhlAn4 provided very good precision at a moderate level of recall, and tunable MAPQ minimum had little effect (Fig. 1g, 2g, S4g, S5g). Sylph delivered moderate to good recall at very high precision, making very few false positive identifications (Fig. 1h, 2h, S4h, S5h). Sylph’s tunable ANI threshold performs best when left at the default of 95% in these datasets. Results were largely congruent between the *Mycobacteriaceae* and *Neisseriaceae* datasets, although overall classifier performance, as measured by AUC, was lower for *Neisseriaceae*. This likely reflected both the overall higher degree of HGT and genome relatedness in this family and the lower absolute number of reads simulated (as simulations were parameterized based on a specified depth, and *Neisseriaceae* have shorter genomes, mean length of 2.4Mb vs 5.7Mb for *Mycobacteriaceae*). PR curves estimated using simulated metagenomes with read length 100bp (Fig. 1,2) were comparable to those when a simulated read length of 150bp was used (Fig. S4,S5), although the longer reads led to lower optimal read thresholds, and slightly higher AUC values (Table 1 vs S6). Given the slightly lower AUC values observed with 100bp read lengths, subsequent detailed analyses were performed using primarily the 100bp read sets, as this should permit conservative interpretation of the results. PRC curves, and the F_1_-score were used to define optimal thresholding for each dataset (*Neisseriaceae*, *Mycobacteriaceae*) and method for further testing of recall and precision, below (Table 1, S6).

**Figure 1.**
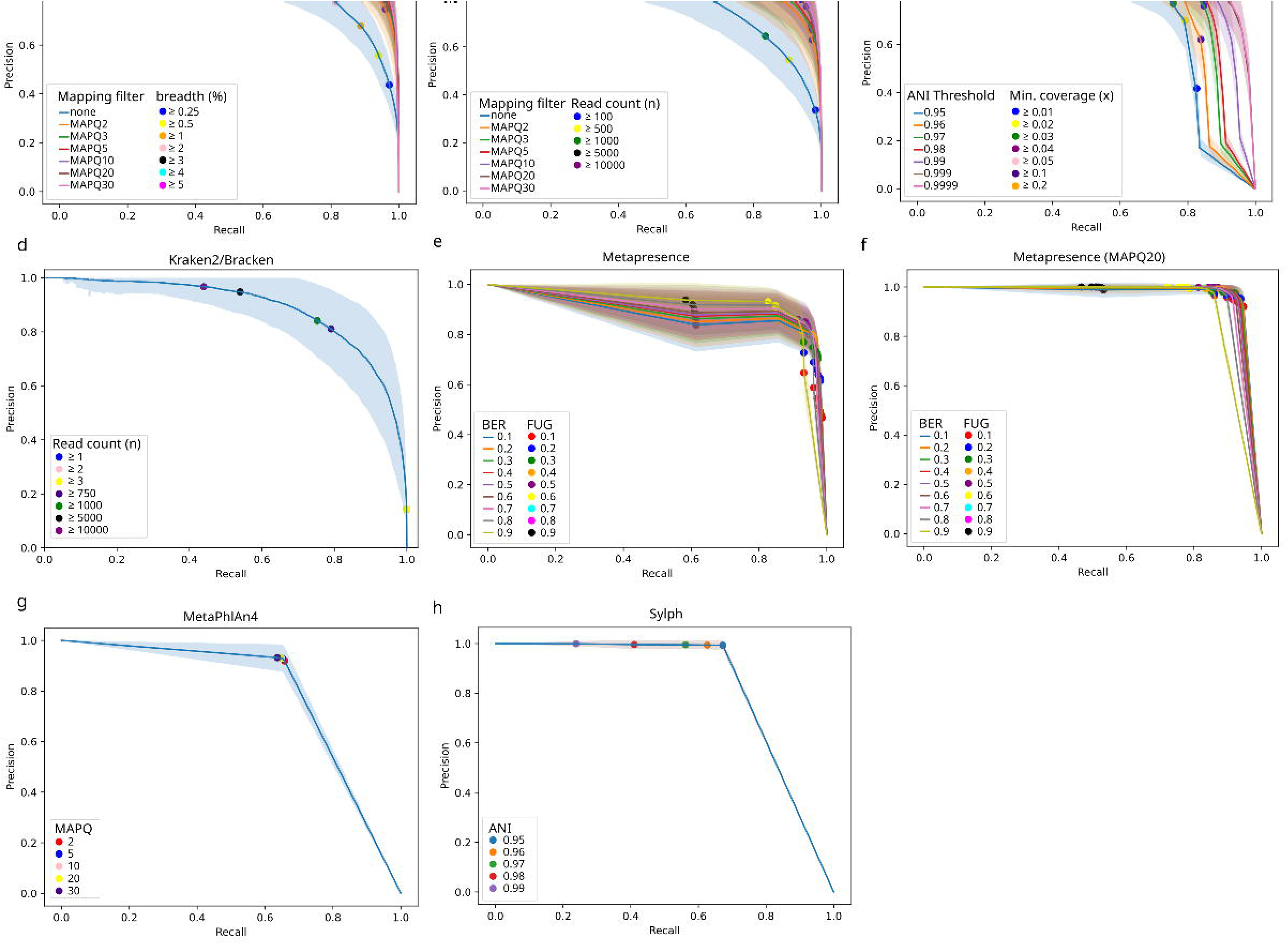
Precision-recall curves (PRCs) for each method, calculated across 500 simulated metagenomes (100bp read length)of *Mycobacteriaceae*. Each panel illustrates the trade-off between precision and recall for each method, PRCs are plotted at the mean precision and recall across replicates, with the standard deviation across replicates shaded. Each panel includes a sample of specific thresholds used to define the curves. Visualized methods are simple aligned breadth of coverage (a) and read count (b) threshold-based methods, both unfiltered and filtered for mapping quality (MAPQ2,3,5,10,20,30). YACHT (c) executed using five different average nucleotide identity (ANI) thresholds, varying the threshold of minimum coverage; Kraken2/bracken (d), executed using a custom index, varying assigned read-count thresholds. Metapresence, executed on the unfiltered read alignment (e). Fraction of Unexpected Gaps (FUG) and Breadth-Expected breadth Ratio (BER) refer to thresholding parameters used by metapresence. Metapresence, executed on the read alignment, filtered for mapping quality at MAPQ20 (f). MetaPhlAn4 (g) executed varying levels of minimum MAPQ filtering, using the default database. Sylph (h), executed with default parameters, varying the ANI parameter.

**Figure 2.**
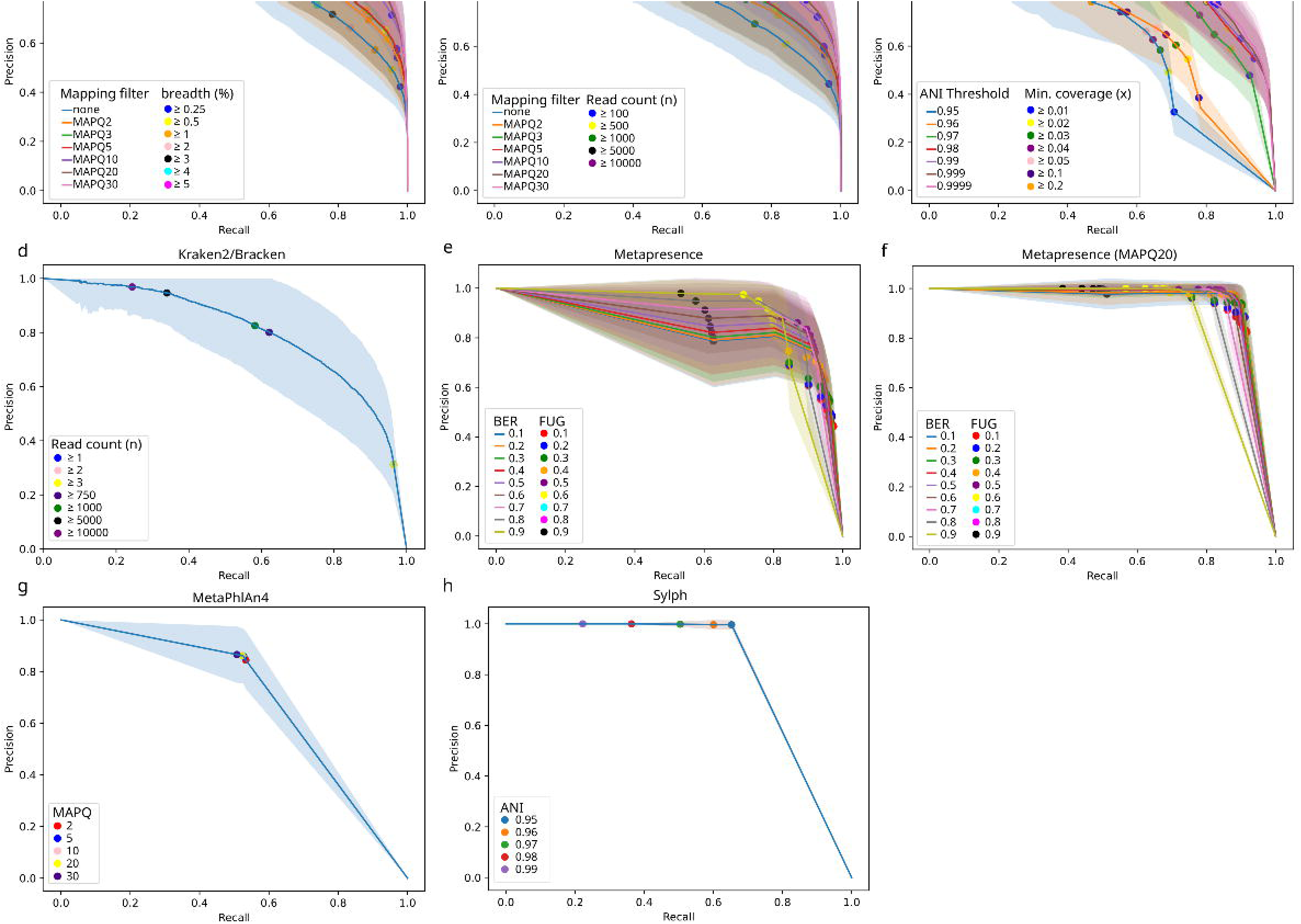
Precision-recall curves (PRCs) for each method, calculated across 500 simulated metagenomes (100bp read length) of Neisseriaceae. Each panel illustrates the trade-off between precision and recall for each method, PRCs are plotted at the mean precision and recall across replicates, with the standard deviation across replicates shaded. Each panel includes a sample of specific thresholds used to define the curves. Visualized methods are simple aligned breadth of coverage (a) and read count (b) threshold-based methods, both unfiltered and filtered for mapping quality (MAPQ2,3,5,10,20,30); YACHT (c) executed using five different average nucleotide identity (ANI) thresholds, varying the threshold of minimum coverage; Kraken2/bracken (d), executed using a custom index, varying assigned read-count thresholds. Metapresence, executed on the unfiltered read alignment (e). Fraction of Unexpected Gaps (FUG) and Breadth-Expected breadth Ratio (BER) refer to thresholding parameters used by metapresence. Metapresence, executed on the read alignment, filtered for mapping quality at MAPQ20 (f). MetaPhlAn4 (g) executed varying levels of minimum MAPQ filtering, using the default database. Sylph (h), executed with default parameters, varying the ANI parameter.

### Recall performance in relation to depth, abundance and accuracy of relative abundance estimates

For all methods, overall recall improved with increasing simulated dataset depth and absolute read abundance, reaching close to perfect recall above depth > 0.1x for all methods with the exception of MetaPhlAn4 (Fig. 3a,b,d,e). MetaPhlAn4 uses species-level genome bins (SGB), which makes direct species-level comparisons imperfect, and unfairly penalizes the method in this analysis, as seen in previous studies^5^. Recall for the breadth-based threshold method varied with alignment quality filtering, related to the optimized threshold chosen (Table 1). Notably, sylph and to a lesser extent MetaPhlAn4 had lower recall at low read abundance, with poor recall at depths of coverage less than 0.1x, which likely explains the lower maximal recall observed in the PR curves, as our simulated datasets had low overall median depth (0.22-0.23x). Recall for Kraken2/bracken was as expected for the optimized read count thresholds used (Table 1). Metapresence had a much lower limit of detection, and was able to identify the presence of genomes from as little as hundreds of reads (Fig. 3b,e). YACHT’s recall performance at low abundance is intermediate between sylph and metapresence at the optimized thresholds (Fig. 3a,b,d,e, Table 1). The recall performance of metapresence demonstrated an unexpected decrease in recall at high depths of coverage, which is suspected to be related to alignment filtering and thresholding parameters (BER/FUG), as the algorithm switches between consideration of both BER and FUG to only BER at higher read depths^6^.

**Figure 3.**
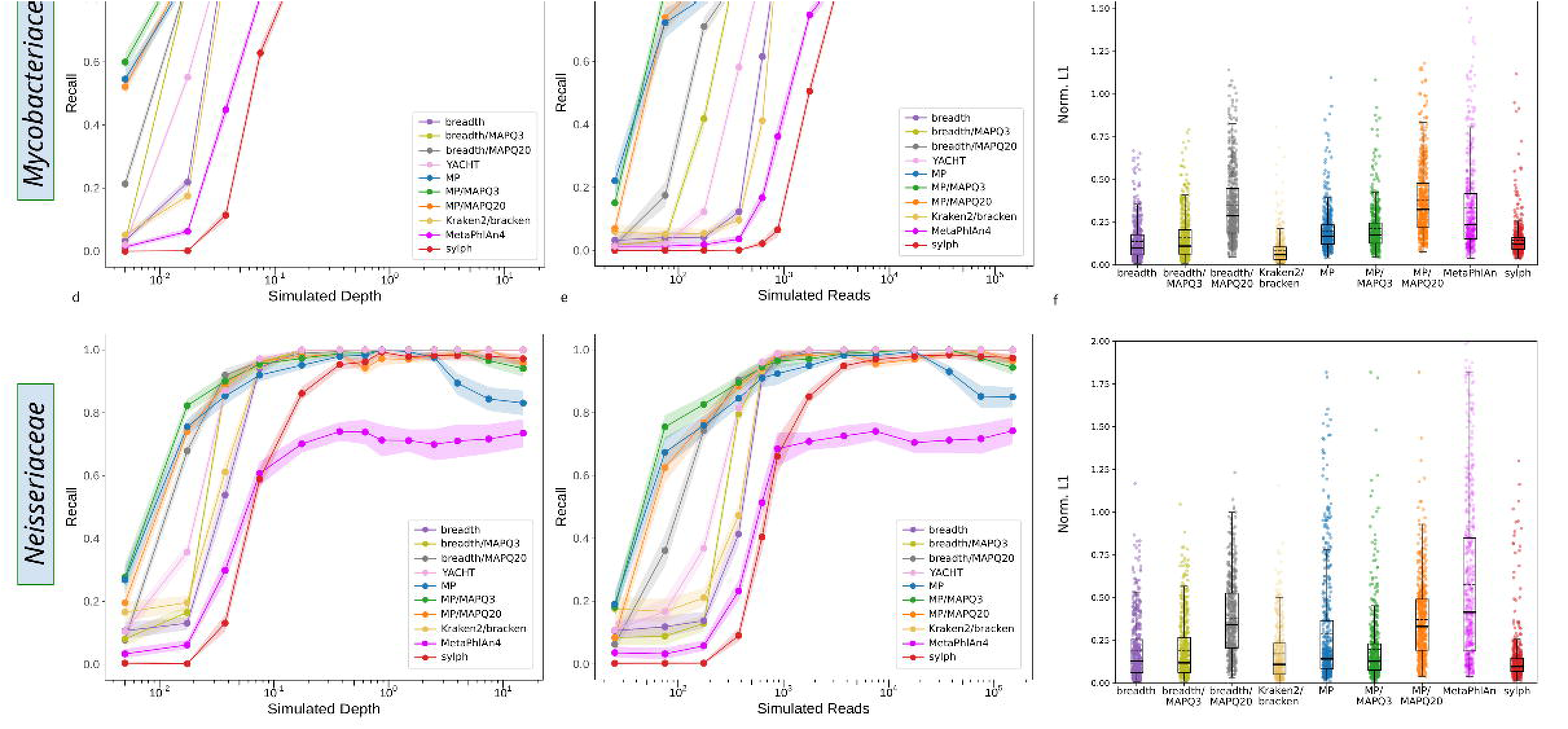
Recall plotted against binned read depth (a,d) used to simulate metagenomic reads as well as the actual number of simulated reads generated (b,e). The x-axis is log-transformed and mean recall is given at the bin midpoint. Results are depicted for the (a,b) *Mycobacteriaceae* and (d,e) *Neisseriaceae* datasets. Recall is summarized in each depth bin (bin boundaries: 0, 0.01x, 0.025x, 0.05x, 0.1x, 0.25x, 0.5x, 0.75x, 1.0x, 2.0x, 3.0x, 5.0x, 10.0x, 10.0x-max) and read count bin (bin boundaries: 0, 50, 100, 250, 750, 1000, 2500, 5000, 10000, 25000, 50000, 100000, 100000-max) across replicates of the simulated metagenomes on a per genome and method basis. The mean recall for the last bin in each plot is placed at an arbitrary x value within the last bin (as the upper border is infinite). Confidence intervals around each method’s estimated recall were computed by a bootstrap resampling procedure using 1000 replicates. Accuracy of relative abundance estimates of the different methods are summarized in (c,f) using the normalized L_1_ metric, with median and interquartile range depicted using box plots, and the mean normalized L1 shown at a dashed line.

Further, at very high read depth > 2.0x, the expected BER against which the BER is compared becomes stricter while calculated BER saturates, which likely explains the decrease in recall observed at high simulated depth. When executed with default rather than optimized BER and FUG thresholds, these dynamics are exaggerated (Fig. S8). With stricter alignment filtering (MAPQ20), metapresence achieved slightly lower recall at higher abundance relative to other methods, particularly with default thresholds (Fig. S8). Also unexpectedly, moderate alignment quality (MAPQ3) filtering increased rather than decreased recall at low abundance (Fig. 3a,d, Fig. S8), which highlights the complex interplay between alignment mapping quality filtering, BER/FUG thresholds, and read spatial distribution: moderately strict filtering decreases overall aligned read count, but in the case of a genome truly present, may make the read distribution more even, thus increasing calculated BER and FUG^6^.

The accuracy of each method to recapitulate the original relative abundance of the individual species in the replicate metagenomes was compared using the normalized L_1_ distance (Fig. 3c,f; Table S7). YACHT does not directly report relative abundance, and so was not included. Sylph, Kraken2/bracken and the breadth of coverage threshold method were the most accurate methods, with low mean normalized L_1_ of between 0.08 and 0.17 for both the *Mycobacteriaceae* and *Neisseriaceae* datasets, while metapresence was slightly less accurate with 0.20 and 0.29, respectively. For all alignment-based methods there was a clear trend toward less accurate abundance estimates at higher alignment quality filtering levels, particularly at MAPQ20 (Table S7). MetaPhlAn4 was the least accurate method.

### Precision performance in the presence of other genomes and incomplete host read removal

Inclusion or not of reads from the CAMI II mock communities affected precision for all methods, although to a varying degree (Fig. 4a,b,d,e, Table S8). Inclusion of CAMI II reads led to a relatively larger decrease in precision in the *Neisseriaceae* relative to the *Mycobacteriaceae* analysis (Fig. 4a vs d). Kraken2/bracken, when executed with an index composed only of our reference genome database, demonstrated very poor precision, consistent with previous estimates^5^, which improved considerably when the full index was used, although it remained poor (Fig. 4a,b,d,e). Sylph and metapresence, with an alignment quality filter, were the best performing methods in terms of precision. Reads derived from more distantly related bacterial genomes simulated from the GTDB had little effect on the methods’ precision, other than for Kraken2/bracken (Fig. 4a,b,d,e). Execution of the methods with the presence of host reads revealed little impact on the precision for these methods (Fig. S10; Table S8).

**Figure 4.**
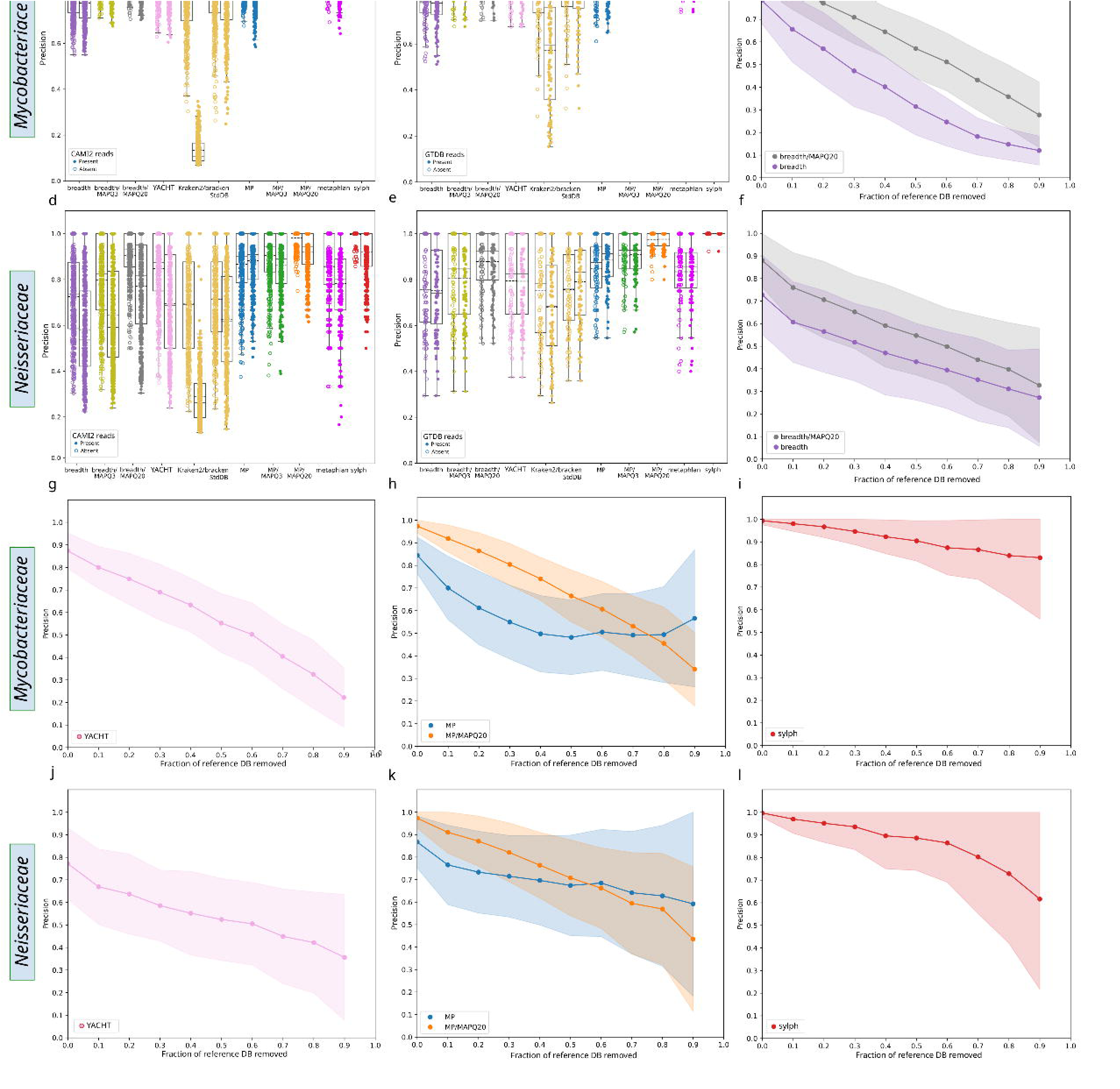
Precision of the examined methods in relation to the presence of reads outside the group of interest (a,b,d,e) and as a function of reference genome database subsampling (c,f,g-l), across the two datasets examined: *Mycobacteriaceae* (a-c, g-i) and *Neisseriaceae* (d-f, j-l). Precision in the face of other microbial reads is calculated both with and without the addition of mock communities from the CAMI II dataset (a,d) and simulated genomic reads derived from 100 randomly sampled genomes from the Genome Taxonomy (GTDB) database (b,e). In both cases, data points represent individual replicates, and boxplots the median and interquartile range. Performance of the methods in the face of an incomplete reference database is shown by subsampling the reference database and recalculating precision at each level of subsampling for the breadth based-classifier (c,f), YACHT (g,j), metapresence (h,k) and sylph (i,l). For the subsampling analysis, points are plotted at mean precision across replicates, and the shaded area represents the standard deviation of precision across replicates.

### Precision performance in the setting of an incomplete reference database

Precision of the breadth threshold method decreased markedly with reference database subsampling (Fig. 4c,f), with alignment quality filtering increasing precision overall but not altering the trajectory of precision loss with a sparse reference genome database. The *Mycobacteriaceae* dataset showed higher overall precision, which may relate to the larger size of the reference database. YACHT demonstrated similar dynamics (Fig. 4g,j). Metapresence without MAPQ filtering as well as sylph showed greater robustness of precision to reference genome database subsampling (Fig. 4h,i,k,l), with metapresence affected to a greater degree than sylph (Fig. 4g,h,i,l). Sylph is relatively robust to the removal of up to approximately 50% of the reference genome sequences before a drop in mean precision below 0.9 was observed, but variation in the range of precision estimates obtained at high subsampling fractions included 1.0 (Fig. 4i,l). Metapresence demonstrated decreased precision after 20-30% of reference genome removal. Unexpectedly, as the subsampling fraction increased, more stringent alignment quality filtering (MAPQ) had a greater relative decay in precision in both datasets (Fig. 4. h,k), similar to YACHT, which likely results from an interaction between the BER/FUG thresholds optimized using the full database and the alignment quality filtering and is less marked when the default, unoptimized BER/FUG thresholds are used (Fig. S9). Variability around estimates are wide, likely reflecting an effect of sampling bias at very low subsampling fractions.

### Case studies

We reanalyzed the Dowdell et al.^10^ dataset of shotgun metagenomic reads characterizing the mycobacterial community in a chloraminated water distribution system (Fig. 5, S11) and examined the distribution of *Neisseriaceae* in the human oral cavity using sequencing data from the human microbiome project^25^ (Fig. 6, S12). We elected to use the two methods, sylph and metapresence/MAPQ20, with the best overall performance in terms of recall, precision and robustness to reference database subsampling (see also discussion).

**Figure 5.**
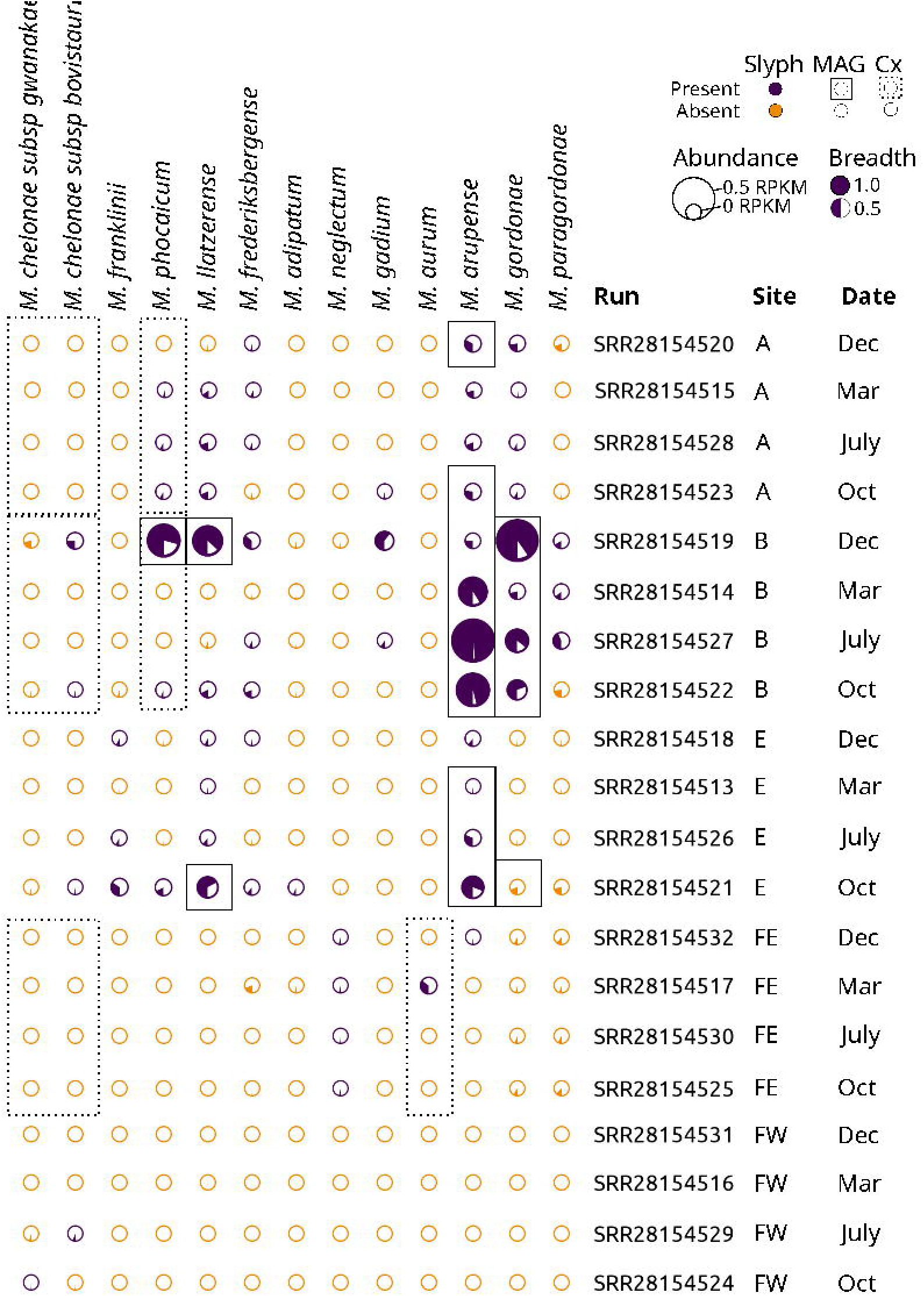
Presence/absence, abundance, and breadth of genomic coverage of mycobacterial species in the metagenomes of a chloraminated drinking water system by the sylph, culture and metagenome-assembled genome (MAG) approaches. This plot relates genomes, ordered phylogenetically as columns, to their estimated presence and abundance in the metagenomes of Dowdell et al^10^. Only mycobacterial species for which any method has been identified are shown. The small circles are shaded according to the breadth of genomic coverage based on unfiltered read alignments and are scaled in size by the abundance in reads per kilobase-million (RPKM), based on the relative abundance reported by sylph (-u option) and scaled by the total read count. Circles colored in purple are identified as present by the sylph method. Those identified by the MAG approach used by Dowdell et al., are identified by a solid square around the circular breadth/abundance plots, while those identified by culture-based methods by Dowdell et al. (pooled by site) are identified by the dashed squares.

**Figure 6.**
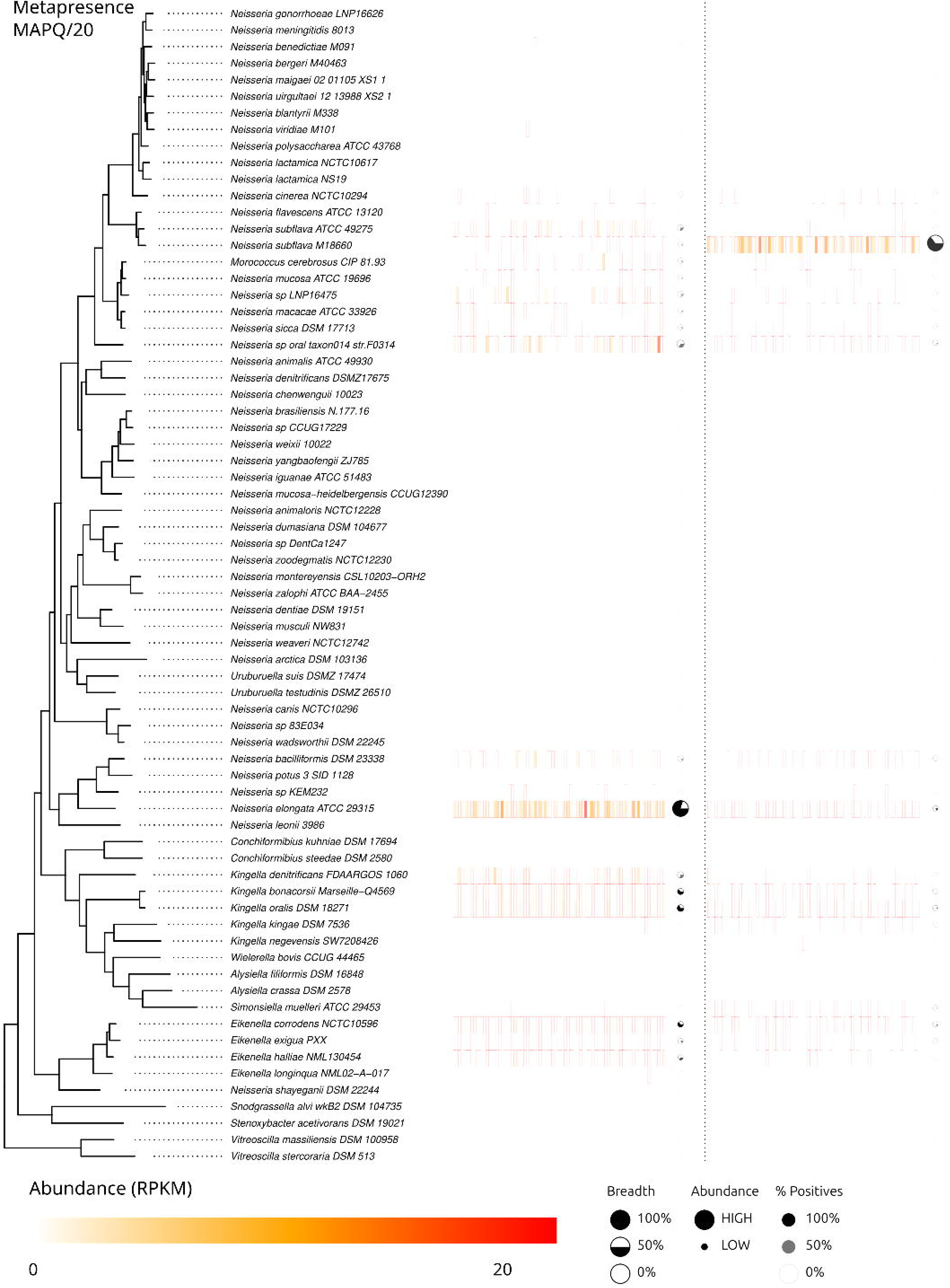
Presence/absence, abundance and breadth of genomic coverage of *Neisseriaceae* species in metagenomic samples from two sites in the oral cavities of humans according to the metapresence method with MAPQ20 alignment filtering. Samples from the Human Microbiome Project representing healthy individuals across two sites: the dorsum of the tongue (left) and supragingival plaque (right). NCBI sequence read archive (SRA) sample accession numbers are provided in Table S5. All *Neisseriaceae* species included in Chenal et al. are shown along with their phylogenetic relationships estimated by core genome analysis^26^. The heatmap represents microbial abundance, scaled to reads per kilobase-million (RPKM), based on the MA filtered alignment. Heatmap cells outlined in red are considered present at a non-zero abundance by metapresence. Circles summarize the data for each species-level taxon across samples, with the size of the circle representing mean abundance, the fraction of the circle filled represents the mean breadth of coverage across samples, and the transparency of the circle represents the number of samples for which the given taxon was detected.

### Mycobacterial communities in a drinking water system

Relative to the approach adopted by Dowdell et al., both sylph (Fig. 5) and metapresence/MAPQ20 (Fig. S11) detected the presence of more genomes of *Mycobacteriaceae* in these samples. This includes those identified by pooled culture-based methods, but not by MAG (e.g. *M. franklinii*, *M. aurum*, and *M. chelonae*). As expected, genomes with high breadth of genomic coverage and abundance are identifiable in both approaches. Genomes estimated to be present at relatively low abundance (e.g. *M. gadium*) were not recovered as high quality MAGs by Dowdell et al., but were detected by sylph and metapresence/MAPQ. Despite the high theoretical sensitivity of culture for mycobacteria, in practice cultivability depends on many factors, and standard approaches may not be optimal for specific mycobacteria^9^. As predicted by the in silico simulations, at low abundance, metapresence with MAPQ20 alignment mapping filter identifies (Fig. S11) the presence of more low-abundance species relative to sylph (Fig. 5). Dowdell et al. also assembled a mycobacterial MAG that they were not able to identify, but this sequence is not available to compare with or augment our reference database, and so may not be reflected in this analysis.

### Presence of specific *Neisseriaceae* in different oral metagenomes

Analysis of the tropism of *Neisseriaceae* in oral metagenomes demonstrates differences in abundance and niche within even the small confines of the human oral cavity (metapresence/MAPQ20 in Fig. 6, and sylph in Fig. S12). *Neisseria subflava* (M18660) is recovered in greater frequency and abundance on the dorsum of the tongue, while *N. elongata* is likewise more consistently detected in the gingival plaque. Multiple species of *Neisseria* known to inhabit the oral mucosa are recovered at lower abundance from both sites. Genera and species associated with teeth/gingiva are recovered more frequently, and at higher abundance, from the supragingival plaque samples (e.g. *N. elongata*, *Eikenella spp.*, *Kingella spp.*). These results are in general congruent for higher abundance taxa with those of Donati et al.^14^ and Albanese and Donati^48^ who used these same HMP data, but employed a different taxonomic database and strain-based approaches: respectively, reference-based alignment followed by SNP calling, as well as MLST profiles^14^ and StrainEst^48^. Relative to these studies, sylph and metapresence detected the presence of more genomes at lower abundance, particularly on the dorsum of tongue. There are differences between the genomes identified by these two methods, particularly when adjudicating the presence of very closely related genomes (e.g. *N. macacae* and *N. sicca*, pairwise ANI=96.5), and in general metapresence/MAPQ20 identifies more species as present (mean 6.6 species per metagenome) relative to sylph (5.7).

## Discussion

Methods to determine the presence of a given genome in a metagenome balance the ability to detect a genome (recall/true positive rate/sensitivity/completeness) against the rate of false positive identifications (precision/purity)^24^. In this use case for these methods, maintaining a high precision is important, and false positive genome identifications are generally to be avoided as it is more conservative to conclude there is lack of evidence to support the presence of a genome than to conclude the incorrect genome is present. By analogy to alternative approaches to identify species of interest - specific PCR or culture are generally relatively precise but sometimes not sensitive (such as when the bacteria is fragile or difficult to grow) or when available PCR primers and the resulting amplicons are a poor match to reference databases or are unable to discriminate to the species level. However, the ideal balance between recall and precision may depend on the exact hypothesis being tested and underscores the importance of a clear understanding of the trade-offs implicit in each method. Fundamentally, there are two sources of information considered by these methods: 1) the degree to which the reads diverge from the reference sequence (ANI, alignment and mapping quality) and 2) the spatial distribution of reads and overall breadth of coverage of reads relative to the reference genome. The datasets studied here, of closely related species in groups with horizontal gene transfers^25,49,50^ and simulated at low overall abundances, incorporating natural genomic variation, likely represent a challenging use case for these algorithms, where consideration of both sources of information is important. This is because HGT may lead to areas of the genome with high pairwise ANI, which may lead to an important number of mapped reads with high ANI, dependent on overall sequencing depth, which may confound some methods. Conversely, presence of closely related genomes may produce many reads that can be mapped throughout a given reference genome, but with lower ANI. This is demonstrated by the overall better performance of the metapresence algorithm and breadth of coverage threshold if the input alignment is filtered for mapping quality (Fig. 1, 2). The focus of this study was performance on species-level taxa, but our dataset also pragmatically includes some reference genomes that are sub-species-level taxa, and so the aggregate performance reflects this.

Overall, this simulation study suggests that the best performing methods for the identification of genomes at the species-level (defined pragmatically), at low abundance and among closely related taxa, are metapresence and sylph. We note that it is important that read alignments are filtered for mapping quality for metapresence (Fig. 1, 2, Table 1, S6), although this has the effect of distorting relative abundance measures (Fig. 3c,f). Therefore, caution is urged in interpreting relative abundance in filtered alignments, which may be used as an initial presence/absence filter prior to recalculation of abundance using an unfiltered alignment.

Simpler alignment threshold-based methods using breadth of coverage and especially read count lack precision in these datasets, particularly if the reference genome database is incomplete (Fig. 4), and enforce thresholds which are dataset specific (Table 1, S6). For a detailed study of a particular taxonomic group, as benchmarked here, these results argue threshold-based methods should employ a simulation-based approach to optimize precision-recall for the specific hypotheses being examined. In a more general study, the default values of BER and FUG for metapresence offer reasonable performance (Table 1; Figs. S8,9) for unfiltered reads, with slightly lower values generally appearing more optimal for MAPQ20 filtered reads. Although difficult to extrapolate precisely, for Kraken2/bracken and alignment-based read count methods, it would seem prudent to adopt a threshold of at least several hundred mapped reads, and closer to 10^3^ when using unfiltered reads to ensure a reasonable degree of precision (Table 1, S6). For methods based on breadth of coverage, a cutoff of between 0.3-0.6% was optimal with quality-filtered reads, while 1.8-3.3% was optimal without filtering, and these may be reasonable starting points for this approach, subject to the caveats above. These cutoffs are also dependent on the read length of the datasets considered. The overall relative performance of methods was similar but in an absolute sense, at a read length of 150bp, several methods had slightly better performance, as reflected by AUC, likely due to the greater specificity of longer reads leading to lower optimal thresholds (Fig. S4, S5, Table S6).

YACHT performed relatively less well in these datasets, which may be related to its ANI thresholding parameter - whereby genomes with pairwise ANI above this threshold are collapsed during sketching^8^, which is particularly problematic for the *Neisseriaceae* database (Fig. S3). Kraken2/bracken, which is widely used for community profiling at higher taxonomic levels, demonstrates important limitations in precision at the species level, particularly in the presence of reads originating from related genomes outside the reference index (Fig. 4a,b,d,e).

These results argue that the detection of species-level genomes by Kraken2/bracken is not reliable^5^ unless conservative thresholds are employed, and this useful tool is better reserved for analyses conducted at the level of genus or higher taxonomic ranks. MetaPhlAn4 employs a marker-gene approach and its overall performance is more similar to that of sylph, but given its use of species-level genome bins, its performance may be underestimated here, as in other studies^5^. Sylph offers high precision and acceptable overall recall, with the lower recall observed here explained by the relatively higher lower limit of detection (here approximately depth 0.1x, Fig. 2a,d) in these datasets characterized by relatively low overall depth. Sylph also accurately recovers the original relative abundances simulated, and is also faster than most other methods^5^, and so would be particularly suited to large scale analyses of thousands of metagenomes, particularly at higher absolute sequencing depths. Sylph is also more robust to the presence of confounding reads outside the group of interest (Fig. 4a,d), and to an incomplete genome database, retaining a reasonable degree of precision even in the absence of up to 50% missing genome sequences from the reference database (Fig. 4i,l). However, the more pronounced decline in precision of these methods observed in the *Neisseriaceae* analysis when CAMI II reads from the ToyHMP dataset are present (Fig. 4d) suggests a possible effect of HGT and/or regions of high genomic similarity among related human commensal organisms outside the reference database and reinforces the need for cautious interpretation of results in these ecosystems. Finally, the declining precision of these methods in the face of an incomplete reference genome database also urges caution in metagenomic studies where a high proportion of uncharacterized taxa or “microbial dark matter” may be present^51^, for example in environmental samples, like soils^9^. For example, an *hsp65* PCR-based study of mycobacterial diversity in global soils found a large majority (> 90%) of amplified sequences were not attributable to known species^9^. This finding suggests a hybrid approach where including as many unique MAGs as possible in the reference database would likely increase the precision of species-level metagenomic profiling, while retaining the added recall at low abundance afforded by a reference-based approach^5^. These conclusions are based on in silico simulations that attempt to capture dynamics of natural genomic variability that lead to genomic divergence of reads from the reference genome within the bounds of real, rather than ideal, species definitions. A limitation of this in silico approach is that dynamics related to relative efficacy of DNA extraction and sequencing between species, among other sampling and wet-lab factors which may bias precision and recall, were not captured.

These results are difficult to compare with those obtained by the authors of the studies describing these methods^5,6,8^, as the methods compared by each author, the nature of datasets used, and simulation procedures all differed somewhat. Among the methods examined here, only metapresence and YACHT have been directly compared^6^. In that study of a mock community generated using an expanded CAMISIM reference genome dataset^52^, the optimal parameters for YACHT were found to be similar to study (ANI = 0.99, min. coverage = 0.05)^6^. Metapresence was found to have higher balanced accuracy and recall (true positive rate) relative to YACHT, particularly when the mock community was created with highly unequal abundance, which is consistent with the greater recall seen here at low sequence abundance (Fig. 3a,b,d,e). Our estimate of sylph’s recall at low abundance and precision is quite similar to those reported by that tool’s authors^5^, where a depth of coverage greater than ∼0.1x is necessary before genomes are reliably detected (Fig. 3a,d) and that precision is generally very high (> 0.9) across a range of conditions (Fig. 1h, 2h, Fig. 4a,b,d,e,i,l).

These approaches should facilitate the study of species-level presence and abundance of a targeted group of species. In the first of two real-world metagenomic datasets examined here, the results of both sylph (Fig. 5) and metapresence (Fig. S11) show that this approach identified more genomes than a MAG-based approach, but concurred when the overall abundance and breadth of coverage was high (Fig. 5). Although this study of mycobacterial communities in a drinking water system did not include an exact gold standard as culture results were pooled, and identification by mass spectrometry may be imprecise^12^, sylph and metapresence detected several species found in culture but not by the MAG approach (e.g. *M. chelonae, M. aurum*), likely because their overall abundance was too low for co-assembly of a high-quality MAG. Both sylph and metapresence/MAPQ20 detected the presence of species, also found in co-assembled MAGs, in more samples relative to the coverM-based approach used by Dowdell et al., although this is likely related to the coverage thresholds used. The importance of consideration of the spatial read distribution and/or read-to-reference ANI/MAPQ is further suggested by the sometimes counterintuitive relationship between read abundance and unfiltered breadth of coverage (illustrated in Fig. 5 and S11), and the determination of presence or absence by these methods, although these relationships were not explored quantitatively in this dataset. This dataset also illustrates some advantages of culture-independent techniques for the profiling of mycobacterial communities^12^, which captured several species (in the case of *M. arupense*, convincingly so), not recovered in culture for a given site. Overall, a more diverse mycobacterial community is suggested by the sylph and metapresence analyses relative to either the MAG or culture-based analysis, including the identification of biologically plausible species recovered from other water systems like *M. gadium*^53,54^ and *M. frederiksbergense*^55^ not detected by the other approaches.

In the second real-world example, the human oral cavity is home to a complex microbiome, where multiple *Neisseriaceae* species not only occupy overlapping niches, but also readily exchange genetic material^25^. The increasing availability of high quality genomic reference sequences in combination with the methods tested here allow rapid delineation of the tropism of *Neisseriaceae* in the oral cavity (Fig. 6, S12), recapitulating the main patterns of tropism seen among high abundance species in strain level methods^14,48^ qualified by important differences in the approach and taxonomic database used. Although the precision of the methods used here is high, it is not perfect in these complex ecosystems, as further demonstrated by the CAMI II analysis (Fig. 4d), and so although the main trends in tropism are evident, finer details of species identification in a given sample at very low abundance among closely related genomes remains challenging, and variable between sylph and metapresence, although with the latter recovering more species at low abundance, as suggested by the benchmarking (Fig. 6 and Fig. S12).

The results of this study demonstrate the feasibility of accurate species-level identification of a defined group of interest within metagenomic samples using these state-of-the-art algorithms. We show that sylph and metapresence (with alignment filtering) can be used for profiling known species in a specific taxonomic group and are robust, to varying degrees, to both incomplete host-sequence removal, and the absence of genomic reference sequences for species outside of the group of interest. Although the focus of our study is not pathogen detection in clinical specimens and this was not directly benchmarked here, the performance of these methods is promising to detect specific pathogenic organisms in complex metagenomic datasets. However, in many cases, pathogenicity can be a matter of strain identity or presence of a specific gene encoding a virulence factor^56^, and so additional analyses beyond simple species (or even sub-species) identification, may be required. Despite their promise, these methods have limitations that shape their ideal use cases. Metapresence with alignment quality filtering has high recall, but suffers from an important loss of precision in the face of a highly incomplete reference, as well as less accurate abundance estimation. For these reasons, metapresence may be well suited to human microbiome and related studies - where high recall in the face of low biomass is advantageous, and the majority of species likely present have been characterized at the genomic level (either by MAG or traditional approaches). For sylph, recall is limited at very low abundance, an important consideration for pathogen detection and low-biomass studies but is balanced by very high precision that is maintained despite a significantly incomplete reference database. This combination of features and the algorithm’s speed would be most advantageous in the face of a highly incomplete reference database and high biomass, as would be expected in, for example in deeply sequenced soil samples.

## Acknowledgments

L.B.H received funding from the Fonds de Recherche du Québec - Santé/Québec Ministère de la Santé et des Services sociaux Clinician-Scientist Training Program. F.J.V. received a Junior 1, Junior 2, and Senior research scholar salary award from the Fonds de Recherche du Québec - Santé. This research was enabled in part by support provided by Calcul Québec and the Digital Research Alliance of Canada (alliancecan.ca). This work was supported by the Natural Sciences and Engineering Research Council of Canada (NSERC) discovery grant (RGPIN-2023-05657) and Canadian Institute of Health Research (CIHR) project grant (450862). Funders had no role in the conception or conduct of the research reported in this manuscript. The authors thank Dr. Marcel Behr for a critical reading of an early draft of this manuscript.

## Data/Code availability

Scripts used to generate the synthetic metagenomes, process the results, and generate the figures are available in a Zenodo repository (10.5281/zenodo.19475466, https://doi.org/10.5281/zenodo.19475466). Truthsets and the results for each analysis are also included in the repository. All genome information used in this analysis are publicly available, with genome accession numbers found in Tables S1, S2, S3, S4, and metagenomic reads for the Dowdell et al. dataset available on NCBI, under Bioproject PRJNA1081894 and those from the Human Microbiome Project also available on NCBI, with SRA accession numbers in Table S5.

## Author Contributions

L.B.H. Conceived the analysis, performed experiments, wrote the initial draft, reviewed and revised the manuscript.

J.O.A. Performed experiments, reviewed and revised the manuscript.

G.C. Performed experiments, reviewed and revised the manuscript.

F.J.V. Conceived the analysis, financial/logistical support, reviewed and revised the manuscript.

