## Supplementary Figures for "Comparative performance of reference-based metagenomic tools to identify species-level taxa among families of bacteria: benchmarking *Mycobacteriaceae* and *Neisseriaceae*"

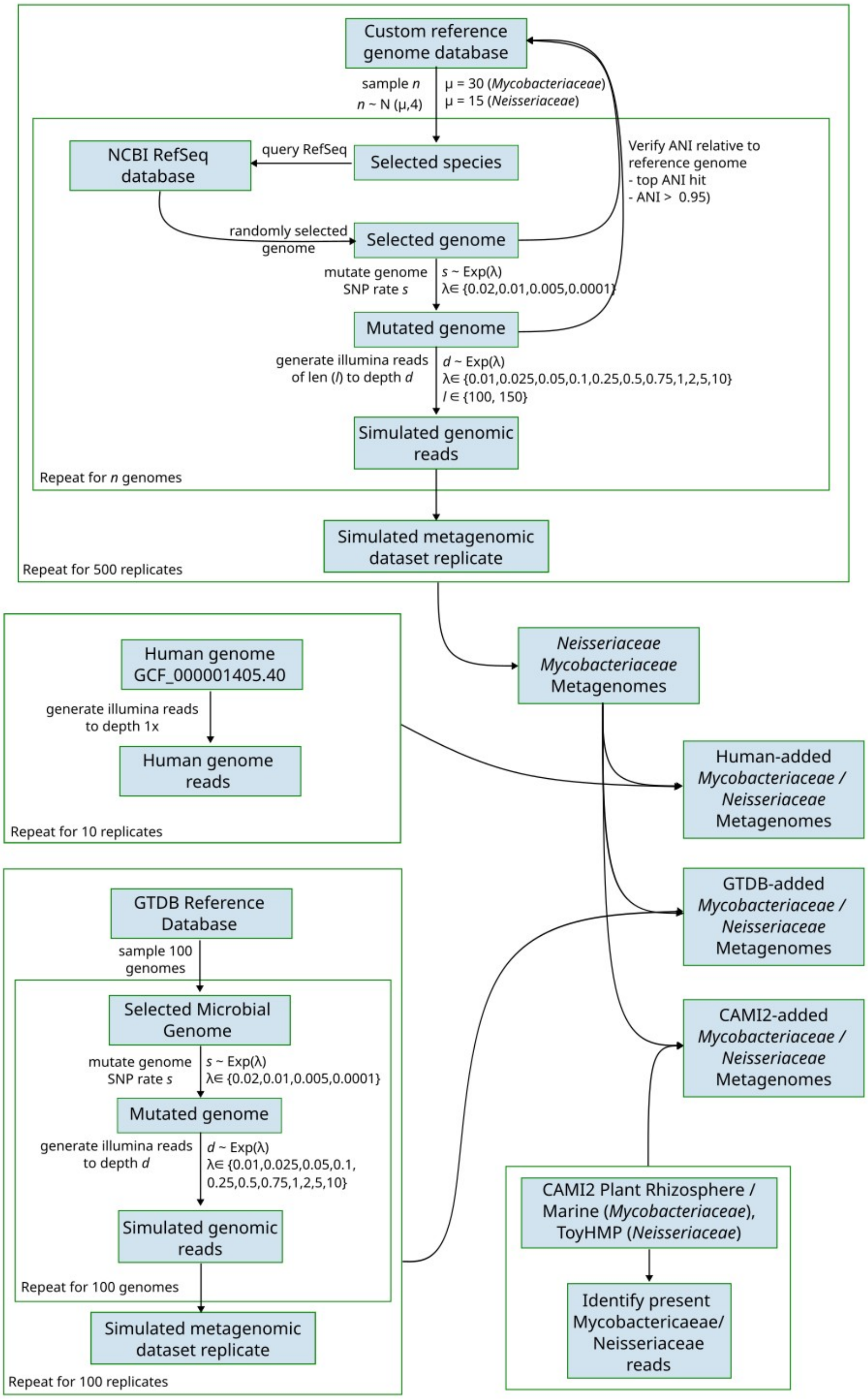

**Supplementary Figure 1.** Diagrammatic summary of the simulation procedure to generate synthetic metagenomes used in this study.

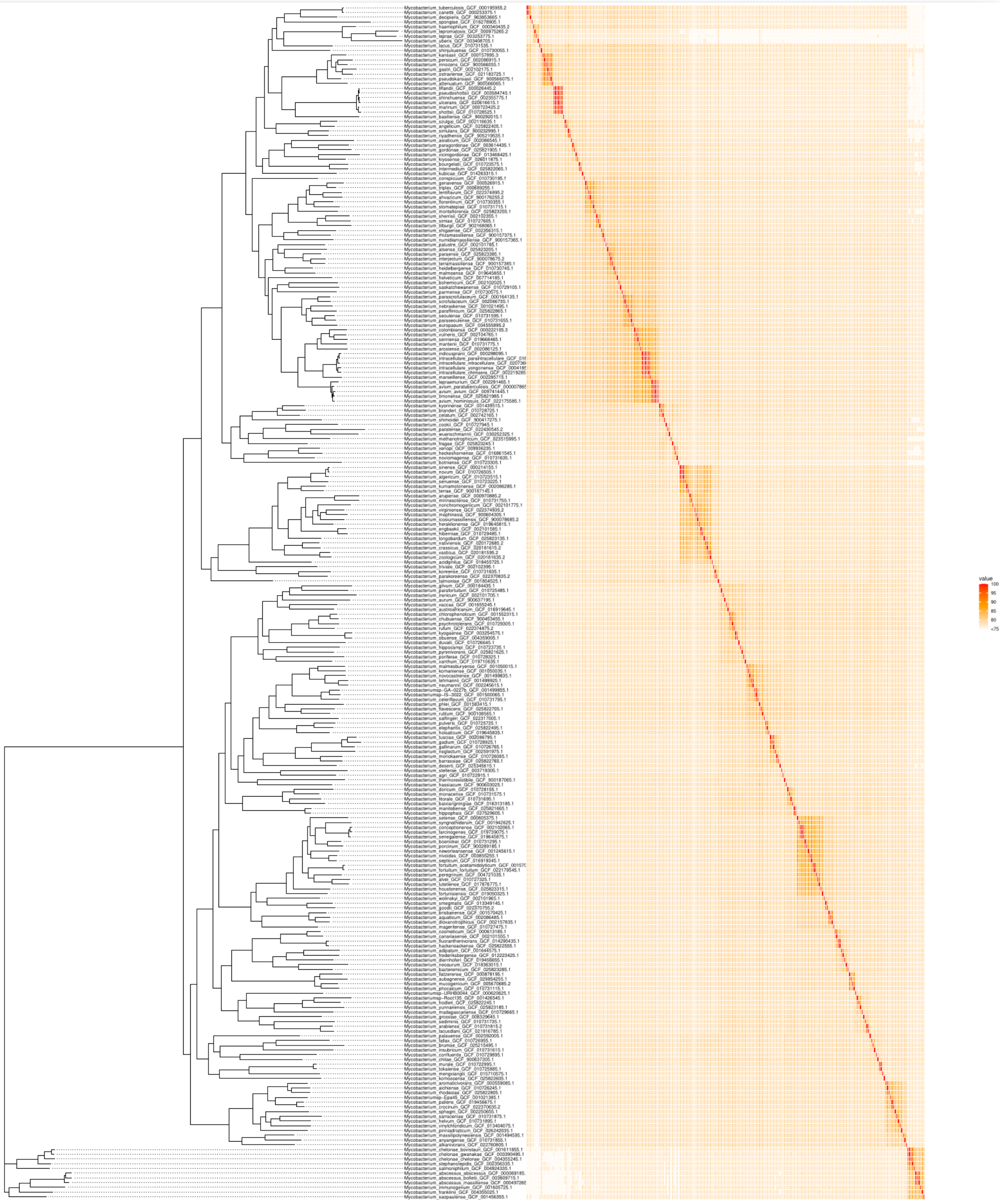
value
100
95
90
85
80
<75

**Supplementary Figure 2.** Core genome phylogeny of the *Mycobacteriaceae* reference database used for read simulation. Pairwise average nucleotide identity (ANI) is illustrated on the right side of the plot.

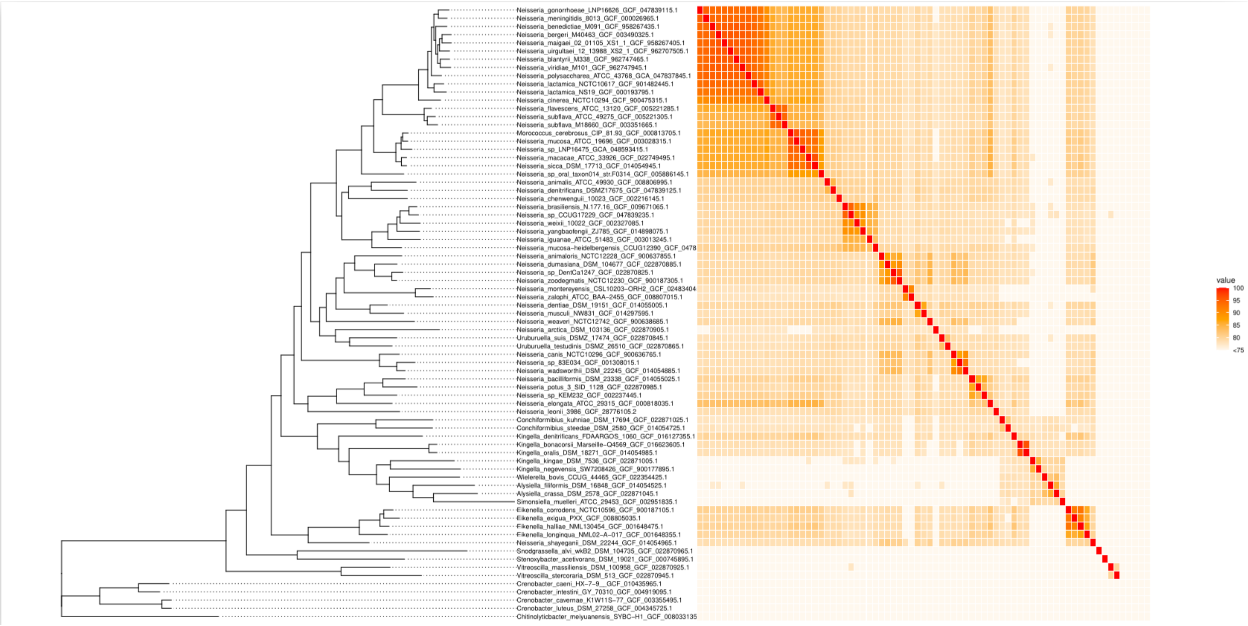


**Supplementary Figure 3.** Core genome phylogeny of the *Neisseriaceae* reference database used for read simulation. Pairwise average nucleotide identity (ANI) is illustrated on the right side of the plot. Several species of *Neisseriales* (*Crenobacter* sp. And *Chitiniloytibacter meiyuanensis*) are included as an outgroup for illustrative purposes but were not used in the analyses presented here.

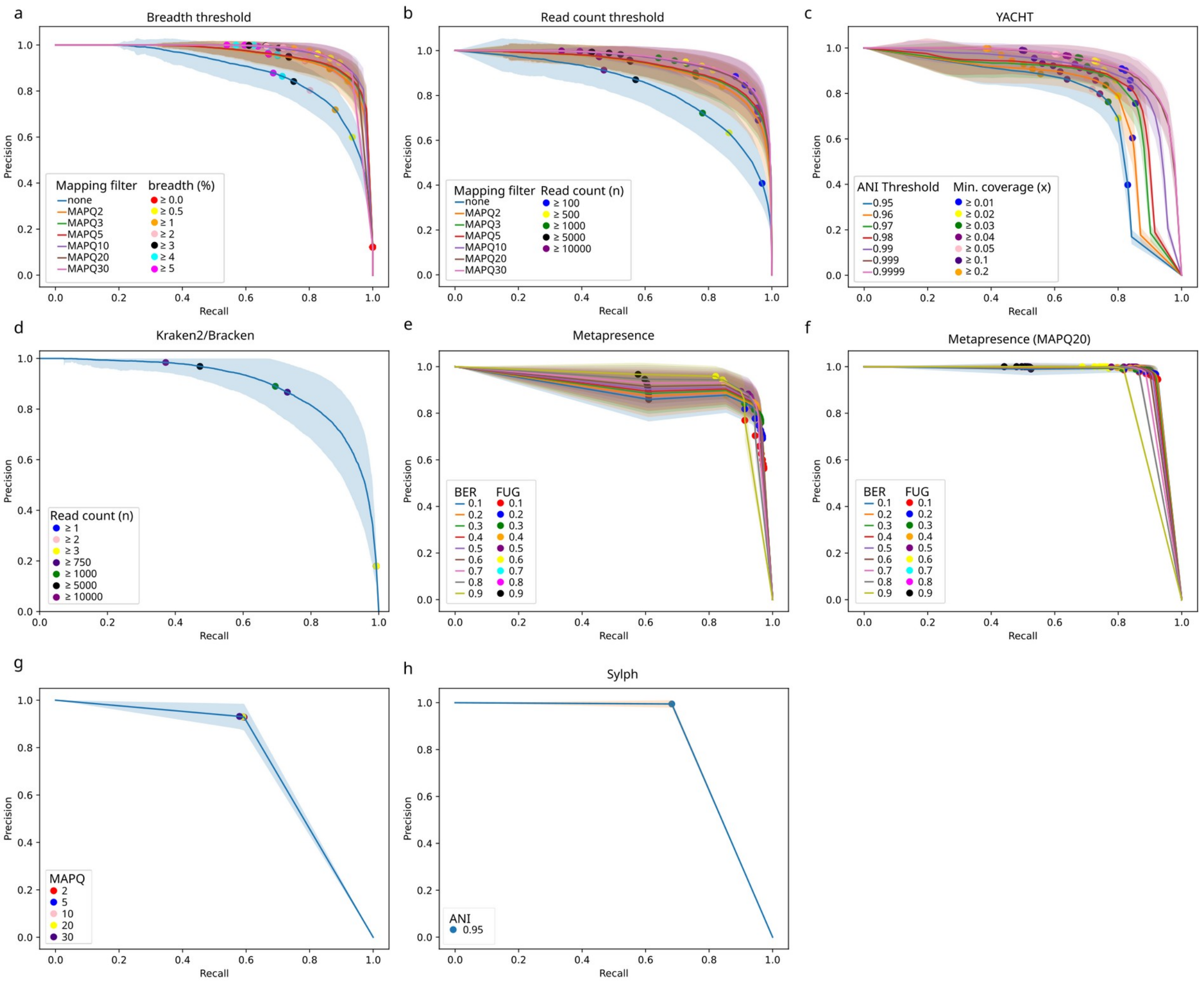


**Supplementary Figure 4.** Precision-recall curves (PRCs) for each method, calculated across 500 simulated metagenomes of *Mycobacteriaceae*using read lengths of 150bp. Each panel illustrates the trade-off between precision and recall for each method, PRCs are plotted at the mean precision and recall across replicates, with the standard deviation across replicates shaded. Each panel includes a sample of specific thresholds used to define the curves. Visualized methods are simple aligned breadth of coverage (a) and read count (b) threshold-based methods, both unfiltered and filtered for mapping quality (MAPQ2,3,5,10,20,30).; YACHT (c) executed using five different average nucleotide identity (ANI) thresholds, varying the threshold of minimum coverage; Kraken2/bracken (d), executed using a custom index, varying assigned read-count thresholds. Metapresence, executed on the unfiltered read alignment (e).

Fraction of Unexpected Gaps (FUG) and Breadth-Expected breadth Ratio (BER) refer to thresholding parameters used by metapresence. Metapresence, executed on the read alignment, filtered for mapping quality at MAPQ20 (f). MetaPhlAn4 (g) executed varying levels of minimum MAPQ filtering, using the default database. Sylph (h), executed with default parameters, varying the ANI parameter.

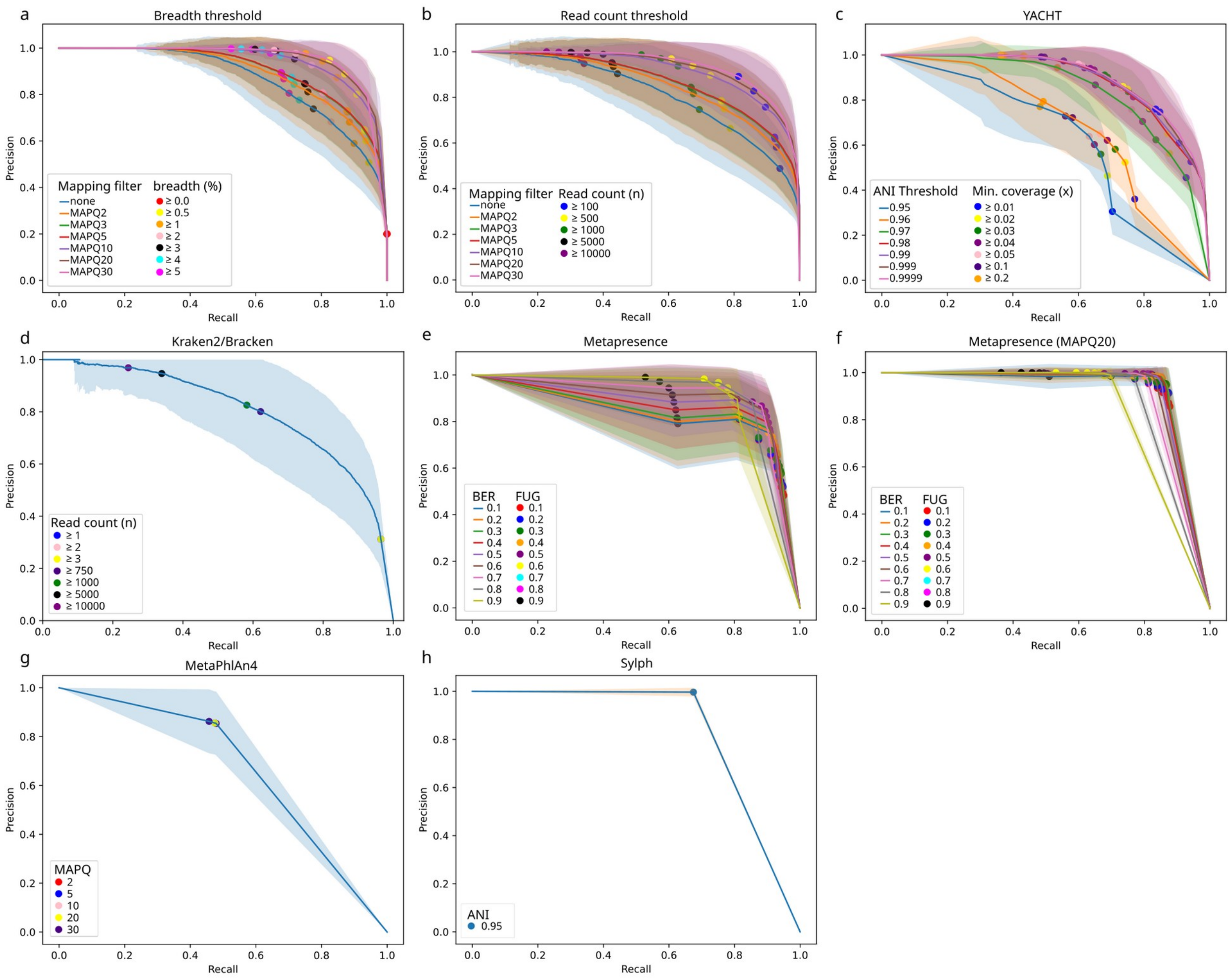


**Supplementary Figure 5.** Precision-recall curves (PRCs) for each method, calculated across 500 simulated metagenomes of *Neisseriaceae* using read lengths of 150bp. Each panel illustrates the trade-off between precision and recall for each method, PRCs are plotted at the mean precision and recall across replicates, with the standard deviation across replicates shaded. Each panel includes a sample of specific thresholds used to define the curves. Visualized methods are simple aligned breadth of coverage (a) and read count (b) threshold-based methods, both unfiltered and filtered for mapping quality (MAPQ2,3,5,10,20,30); YACHT (c) executed using five different average nucleotide identity (ANI) thresholds, varying the threshold of minimum coverage; Kraken2/bracken (d), executed using a custom index, varying assigned read-count thresholds. Metapresence, executed on the unfiltered read alignment (e).

Fraction of Unexpected Gaps (FUG) and Breadth-Expected breadth Ratio (BER) refer to thresholding parameters used by metapresence. Metapresence, executed on the read alignment, filtered for mapping quality at MAPQ20 (f). MetaPhlAn4 (g) executed varying levels of minimum MAPQ filtering, using the default database. Sylph (h), executed with default parameters, varying the ANI parameter.

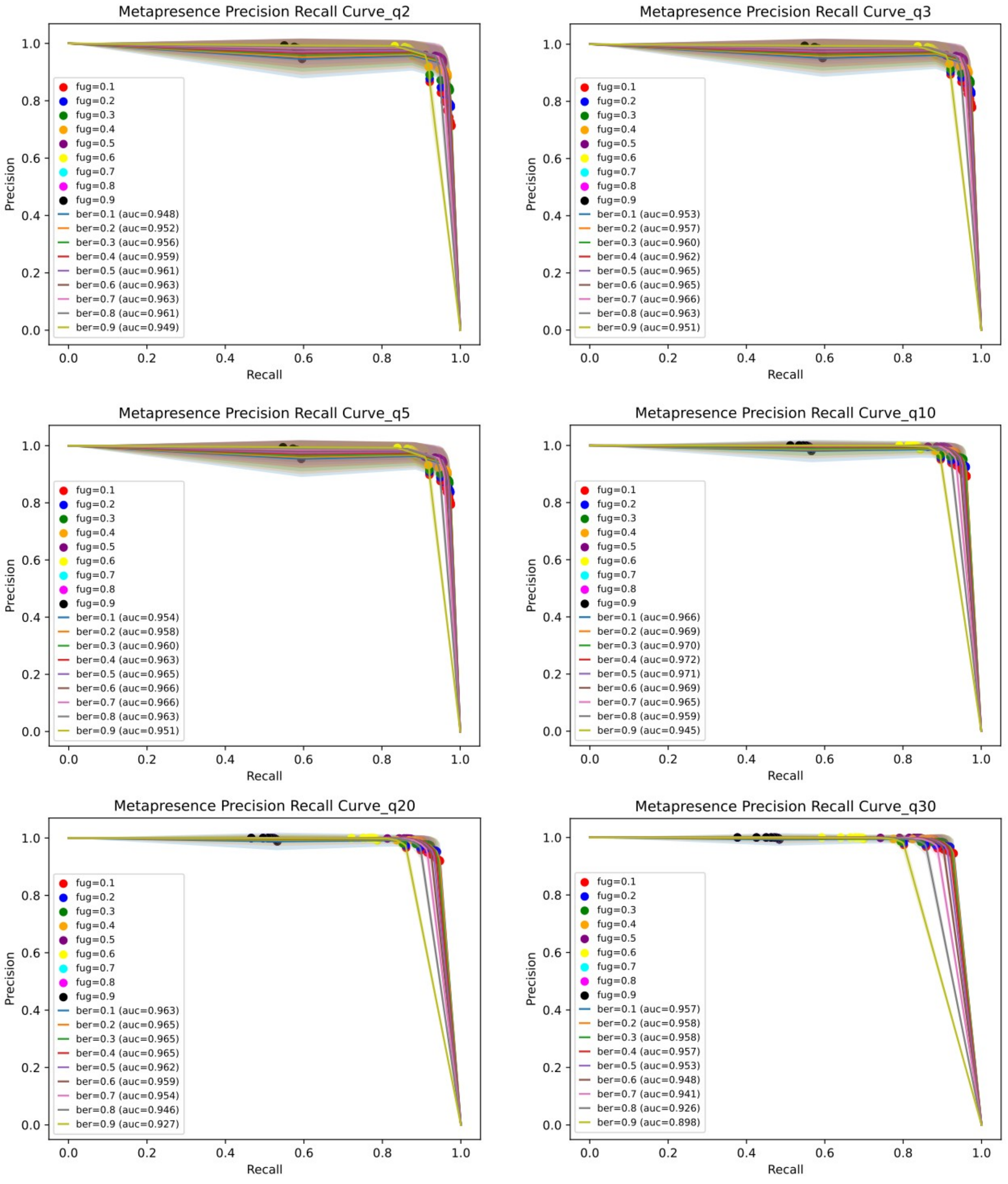


**Supplementary Figure 6:** Additional precision-recall curves for metapresence, at all alignment quality filtering levels (MAPQ 2,3,5,10,20,30) for the *Mycobacteriaceae* dataset.

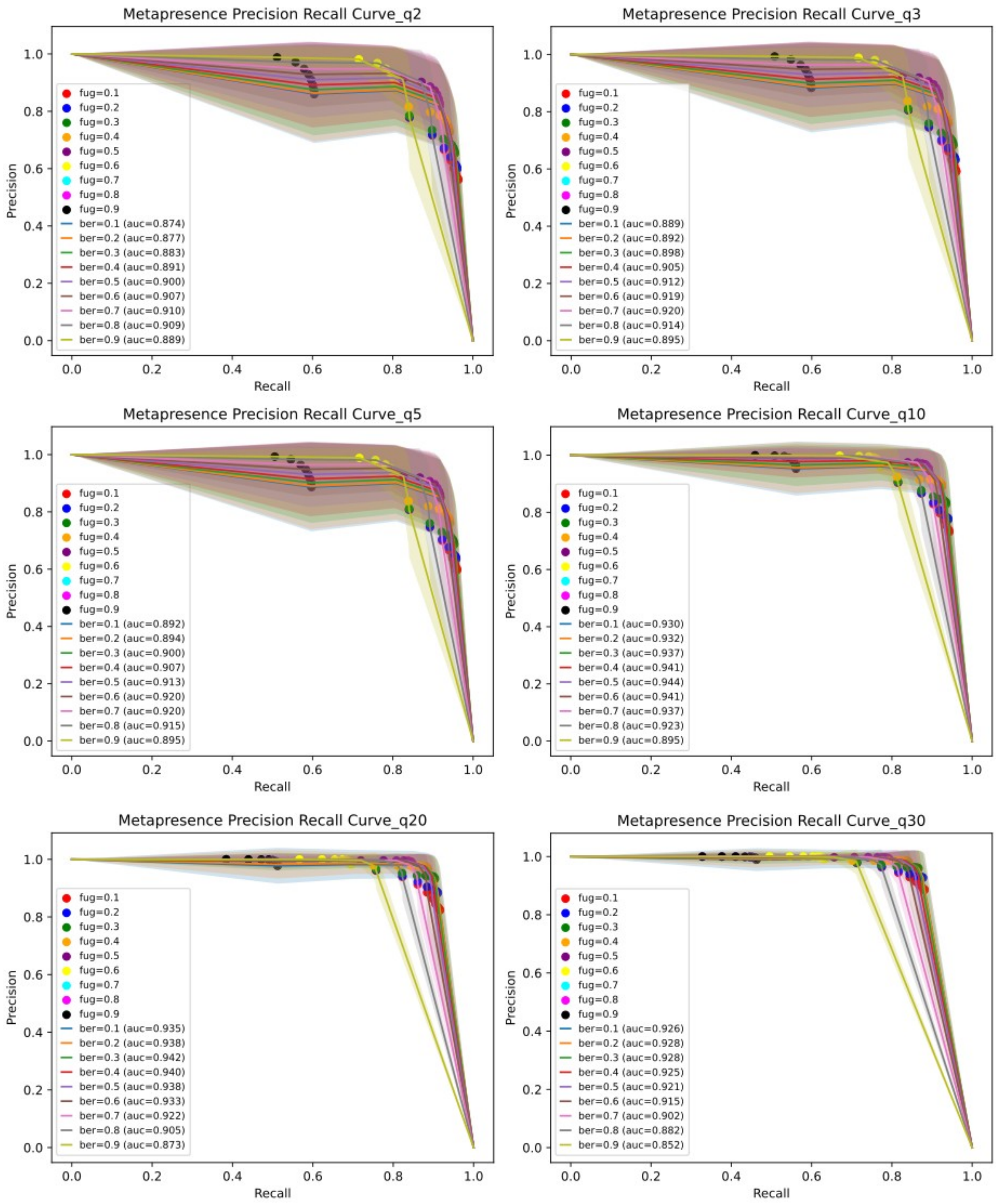


**Supplementary Figure 7:** Additional precision-recall curves for metapresence, at all alignment quality filtering levels (MAPQ 2,3,5,10,20,30) for the *Neisseriaceae* dataset.

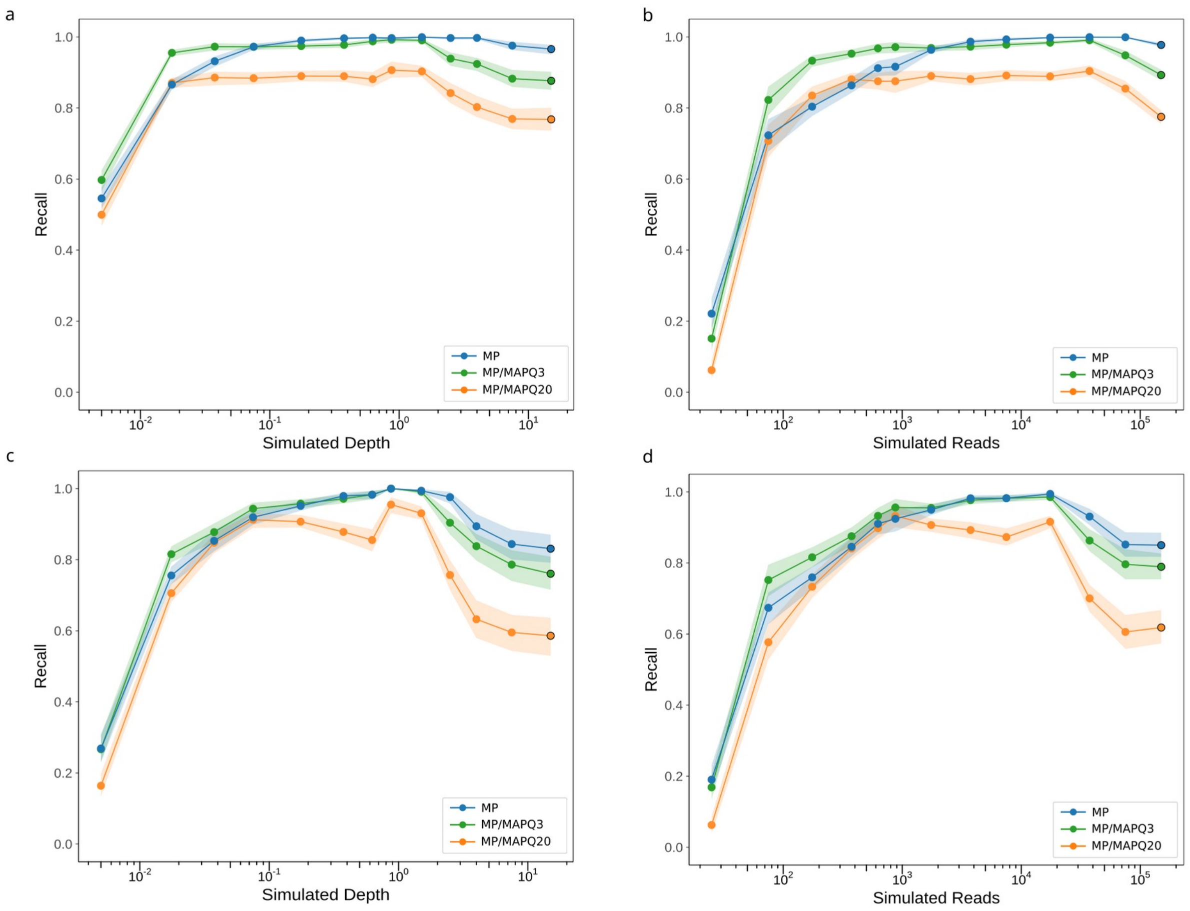


**Supplementary Figure 8:** Recall of the metapresence method as a function of simulated depth (a,c) and simulated reads (b,d) for the *Mycobacteriaceae* (a,b) and *Neisseriaceae* (c,d) simulated datasets. In these plots, metapresence was executed with default thresholds for BER and FUG: 0.8 and 0.5 respectively, rather than optimized values.

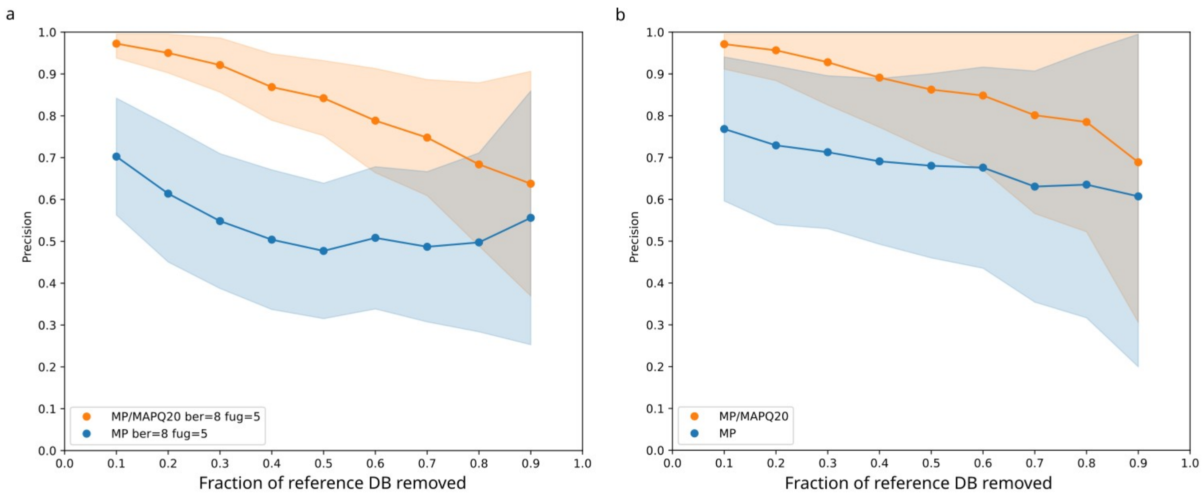


**Supplementary Figure 9:** Precision of the metapresence method plotted against subsampling of the reference genome databases (*Mycobacteriaceae*: a and *Neisseriaceae*: b) using default rather than optimized parameters for the BER and FUG thresholding parameters.

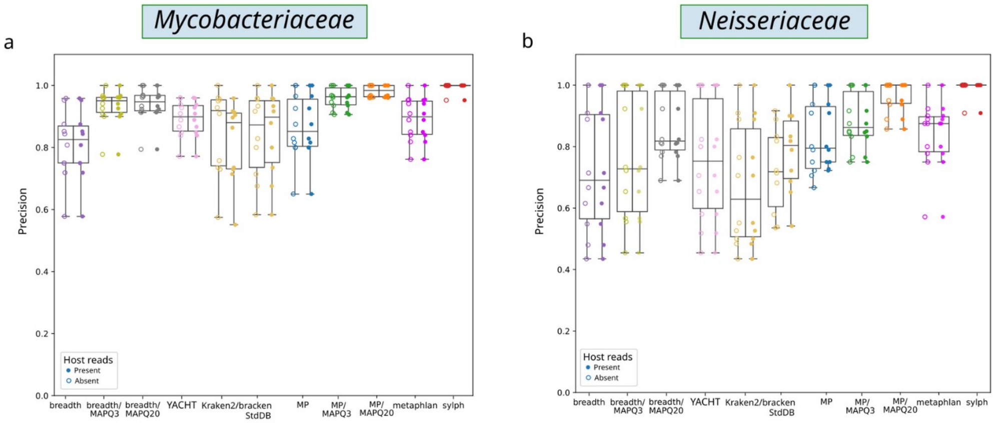


**Supplementary Figure 10.** Precision the face of incomplete host removal calculated both with and without genomic reads from the human genome for both the *Mycobacteriaceae* (a) and *Neisseriaceae* datasets (b).

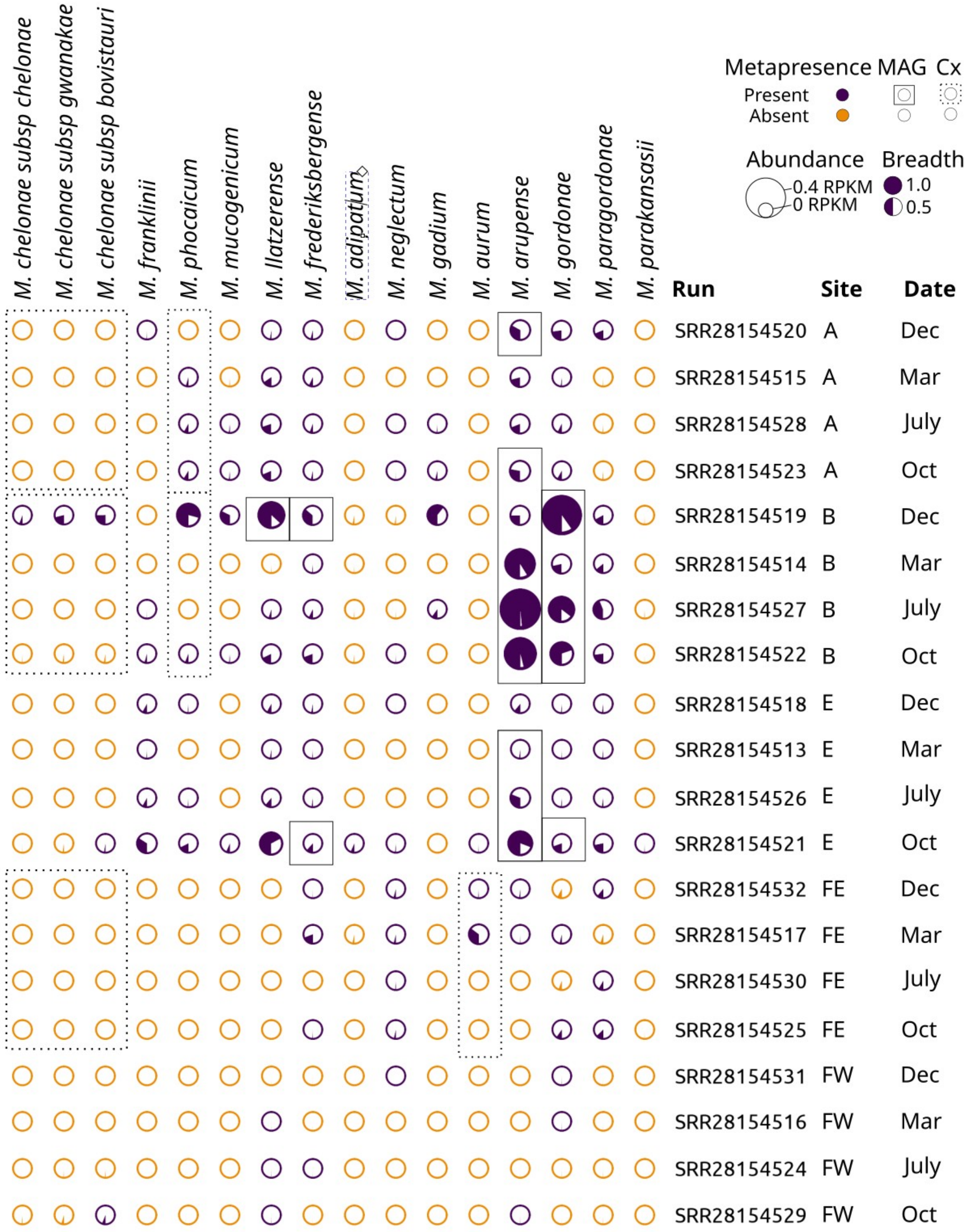


**Supplementary Figure 11:** Presence/absence, abundance, and breadth of genomic coverage of mycobacterial species in the metagenomes of a chloraminated drinking water system by the metapresence with MAPQ alignment filtering compared with different methodological approaches. This plot relates genomes, ordered phylogenetically as columns, to their estimated presence and abundance in the metagenomes of Dowdell et al[10]. Only mycobacterial species for which any method has been identified are shown. The small circles are shaded according to the unfitlered breadth of genomic coverage based on read alignments and are scaled in size by the abundance in reads per kilobase-million (RPKM), based on the read count reported by metapresence/MAPQ20. Circles colored in purple are identified as present by the metapresence method. Those identified by the MAG approach used by Dowdell et al., are identified by a solid square, while those identified by culture-based methods by Dowdell et al. (pooled by site) are identified by the dashed squares.

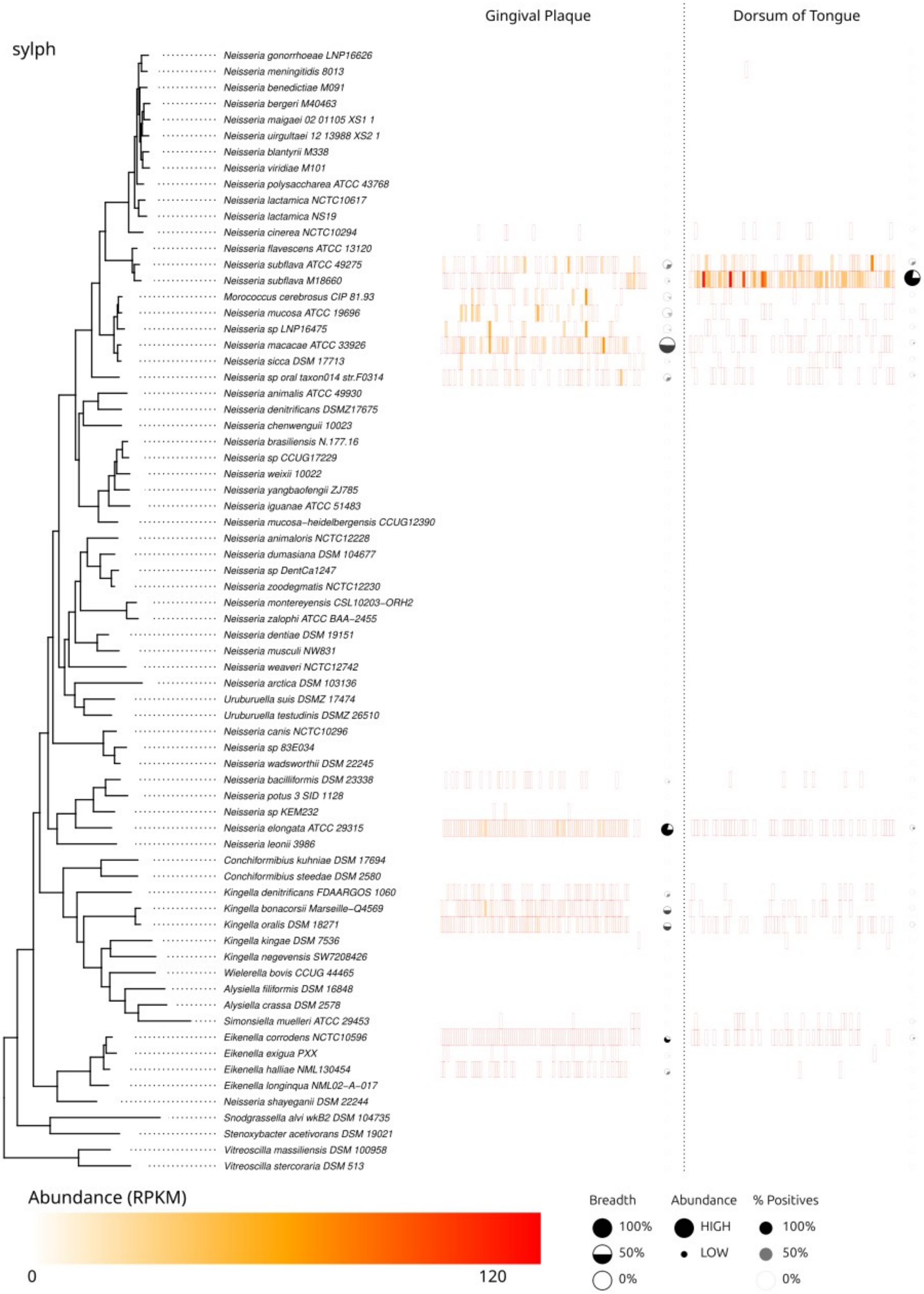

Gingival Plaque
Dorsum of Tongue
sylph
Neisseria gonorrhoeae LNP16626
Neisseria meningitidis 8013
Neisseria benedictiae M091
Neisseria bergeri M40463
Neisseria maigaei 02 01105 XS1 1
Neisseria uirgultaei 12 13988 XS2 1
Neisseria blantyrii M338
Neisseria viridiae M101
Neisseria polysaccharea ATCC 43768
Neisseria lactamica NCTC10617
Neisseria lactamica NS19
Neisseria cinerea NCTC10294
Neisseria flavescens ATCC 13120
Neisseria subflava ATCC 49275
Neisseria subflava M18660
Morococcus cerebrosus CIP 81.93
Neisseria mucosa ATCC 19696
Neisseria sp LNP16475
Neisseria macacae ATCC 33926
Neisseria sicca DSM 17713
Neisseria sp oral taxon014 str.F0314
Neisseria animalis ATCC 49930
Neisseria denitrificans DSMZ17675
Neisseria chenwenguii 10023
Neisseria brasiliensis N.177.16
Neisseria sp CCUG17229
Neisseria weixii 10022
Neisseria yangbaofengii ZJ785
Neisseria iguanae ATCC 51483
Neisseria mucosa-heidelbergensis CCUG12390
Neisseria animaloris NCTC12228
Neisseria dumasiana DSM 104677
Neisseria sp DentCa1247
Neisseria zoodegmatis NCTC12230
Neisseria montereyensis CSL10203-ORH2
Neisseria zalophi ATCC BAA-2455
Neisseria dentiae DSM 19151
Neisseria musculi NW831
Neisseria weaveri NCTC12742
Neisseria arctica DSM 103136
Uruburuella suis DSMZ 17474
Uruburuella testudinis DSMZ 26510
Neisseria canis NCTC10296
Neisseria sp 83E034
Neisseria wadsworthii DSM 22245
Neisseria bacilliformis DSM 23338
Neisseria potus 3 SID 1128
Neisseria sp KEM232
Neisseria elongata ATCC 29315
Neisseria leonii 3986
Conchiformibius kuhniae DSM 17694
Conchiformibius steedae DSM 2580
Kingella denitrificans FDAARGOS 1060
Kingella bonacorsii Marseille-Q4569
Kingella oralis DSM 18271
Kingella kingae DSM 7536
Kingella negevensis SW7208426
Wielerella bovis CCUG 44465
Alysiella filiformis DSM 16848
Alysiella crassa DSM 2578
Simonsiella muelleri ATCC 29453
Eikenella corrodens NCTC10596
Eikenella exigua PXX
Eikenella halliae NML130454
Eikenella longinqua NML02-A-017
Neisseria shayeganii DSM 22244
Snodgrassella alvi wkB2 DSM 104735
Stenoxybacter acetivorans DSM 19021
Vitreoscilla massiliensis DSM 100958
Vitreoscilla stercoraria DSM 513
Abundance (RPKM)
0
120
Breadth
100%
50%
0%
Abundance
HIGH
LOW
% Positives
100%
50%
0%

**Supplementary Figure S12.** Presence/absence, abundance and breadth of genomic coverage of *Neisseriaceae* species in metagenomic samples from two sites in the oral cavities of humans according to the sylph method. Eight samples from the Human Microbiome Project representing healthy individuals across two sites are illustrated: the dorsum of the tongue (left) and supragingival plaque (right). NCBI sequence read archive (SRA) sample accession numbers are provided in Table S5. All *Neisseriaceae* species included in Chenal et al. are shown along with their phylogenetic relationships estimated by core genome analysis[26]. The heatmap represents microbial abundance, scaled to reads per kilobase-million (RPKM), based on the unfiltered alignment (therefore higher absolute abundance relative to Fig. 6). Heatmap cells outlined in red are considered present at a non-zero abundance by sylph. Circles summarize the data for each species-level taxon across samples, with the size of the circle representing mean abundance, the fraction of the circle filled represents the mean breadth of coverage across samples, and the transparency of the circle represents the number of samples for which the given taxon was detected.
